# Translocation of YopJ family effector proteins through the VirB/VirD4 T4SS of *Bartonella*

**DOI:** 10.1101/2024.03.23.586424

**Authors:** Katja Fromm, Monica Ortelli, Alexandra Boegli, Christoph Dehio

## Abstract

The evolutionary conserved YopJ family comprises numerous type-III-secretion system (T3SS) effectors of diverse mammalian and plant pathogens that acetylate host proteins to dampen immune responses. Acetylation is mediated by a central acetyltransferase domain that is flanked by conserved regulatory sequences, while a non-conserved N-terminal extension encodes the T3SS-specific translocation signal. *Bartonella* spp. are facultative-intracellular pathogens causing intra-erythrocytic bacteremia in their mammalian reservoirs and diverse disease manifestations in incidentally infected humans. Bartonellae do not encode a T3SS, but most species possess a type-IV-secretion system (T4SS) to translocate *Bartonella* effector proteins (Beps) into host cells. Here we report that the YopJ homologs present in Bartonellae species represent genuine T4SS effectors. Like YopJ family T3SS effectors of mammalian pathogens, the ‘*Bartonella* YopJ-like effector A’ (ByeA) of *Bartonella taylorii* also targets MAP kinase signaling to dampen pro-inflammatory responses, however, translocation depends on a functional T4SS. A split-NanoLuc luciferase-based translocation assay identified sequences required for T4SS-dependent translocation in conserved regulatory regions at the C-terminus and proximal to the N-terminus of ByeA. The T3SS effectors YopP from *Yersinia enterocolitica* and AvrA from *Salmonella* Typhimurium were also translocated via the *Bartonella* T4SS, while ByeA was not translocated via the *Yersinia* T3SS. Our data suggest that YopJ family T3SS effectors may have evolved from an ancestral T4SS effector, such as ByeA of *Bartonella*. In this evolutionary scenario, the signal for T4SS-dependent translocation encoded by N- and C-terminal sequences remained functional in the derived T3SS effectors due to the essential role these sequences coincidentally play in regulating acetyltransferase activity.

**Significance Statement:** Bacterial pathogens use diverse secretion systems to translocate effector proteins into eukaryotic host cells. Evolutionary successful translocation systems and effector proteins have been acquired by many plant and animal pathogens via horizontal gene transfer. The YopJ family comprises numerous T3SS effectors that share a unique acetyltransferase activity that interferes with various host cell functions. Our study revealed that YopJ homologs in the pathogen *Bartonella* are genuine T4SS effectors and implies an evolutionary scenario in which T3SS-dependent YopJ family effectors may have evolved from such an ancestral T4SS effector by fusion of an N-terminal type-III-secretion signal. Such switches in secretion system specificity of host-targeted effectors may represent an underappreciated phenomenon in effector evolution.

## Introduction

Numerous Gram-negative bacteria utilize either type III (T3SS) or type IV (T4SS) secretion systems to translocate effector proteins into eukaryotic host cells (1, 2) in order to subvert host cellular functions and to downregulate immune responses (3). The YopJ family of T3SS-translocated effectors was named by the prototypic *Yersinia* outer protein J (YopJ) of *Y. pestis* and *Y. pseudotuberculosis* (the closely related *Y. enterocolitica* homolog is known as YopP) and is highly conserved amongst animal and plant pathogens (4). YopJ family effectors share a conserved catalytic triad (histidine-glutamate-cysteine or histidine-aspartate-cysteine) that facilitates acetylation of target proteins (5–8). The acetyltransferase activity is regulated by two cofactors that are confined to eukaryotic host cells: acetyl-CoA and inositol hexakiphosphate (IP6) (8, 9). Binding of IP6 induces a conformational change required for binding of acetyl-CoA which then serves as acetyl donor in the acetylation reaction (8). YopJ/YopP of *Yersinia* acetylate upstream components of mitogen-activated protein kinases (MAPKs) and nuclear factor kappa B (NF-κB) signaling pathways and thereby suppresses innate immune responses, e.g., the secretion of pro-inflammatory cytokines (10–12).

Recently, YopJ homologs were identified in several *Bartonella* species (13–15). Bartonellae are Gram-negative, facultative-intracellular pathogens that are divided into four distinct lineages (16, 17). No effector-translocating T3SS has been identified in any *Bartonella* species (18), thus questioning if and how YopJ homologs may be translocated into host cells.

Noteworthy, all species of *Bartonella* lineages 3 and 4 encode the VirB/VirD4 type-IV-secretion system (T4SS) and translocate *Bartonella* effector proteins (Beps) as essential virulence factors to colonize their specific mammalian host (19–21). The genes encoding the VirB/VirD4 machinery along at least one ancestral Bep were independently acquired by these two lineages via horizontal gene transfer. The subsequent expansion of the Bep repertoires by lineage-specific gene duplication and diversification events increased the capacity to adapt to new mammalian hosts, resulting in rapid speciation by adaptive radiation (22). The *Bartonella* VirB/VirD4 T4SS is composed of VirB2-11 and VirD4 (23, 24). The VirB proteins assemble a large macromolecular translocation channel. Substrates of the T4SS are recruited by the T4SS-associated ATPase VirD4, also termed T4S coupling protein (T4CP). T4SS effectors typically possess a C-terminal secretion signal composed of a few positively charged or hydrophobic residues (25, 26). The secretion signal of the Beps is more complex containing additionally to a positively charged tail sequence an adjacent BID (Bep intracellular delivery) domain of approximately 100 aa (21, 27).

In this study, we report that the *Bartonella* YopJ-like effector A (ByeA) represents a genuine T4SS effector. Utilizing a split-NanoLuc luciferase-based translocation assay, we demonstrate that ByeA translocation via the VirB/VirD4 T4SS depends on N- and C-terminal helices which, as part of the binding sites for IP6 and acetyl-CoA, are broadly conserved regulatory sequences of YopJ family effectors. Consistently, we observed translocation of the YopJ family effectors YopP of *Y. enterocolitica* and AvrA of *Salmonella* Typhimurium via the VirB/VirD4 T4SS of *B. henselae*. In contrast, ByeA was not translocated via the T3SS of *Y. enterocolitica*. The T3SS-specific translocation signal is located in the unstructured, less conserved N-terminal region of YopJ family effectors (28–30). Based on this observation, it is tempting to speculate that YopJ family effectors evolutionary originated as genuine T4SS effectors. Accordingly, acquisition of an N-terminal T3SS signal may then have facilitated the horizontal expansion of this effector family to pathogens harboring T3SS.

## Results

### Genome analysis indicates co-evolution of *Bartonella* YopJ-like effectors with the VirB/VirD4 T4SS in lineages 3 and 4

YopJ homologs were previously identified in several *Bartonella* species, but their biological function and evolutionary history remained elusive (14, 15). Other than their numerous T3SS-translocated homologs in the YopJ family, these *Bartonella* YopJ-like effectors (Bye) cannot be translocated via a T3SS system, as no dedicated-effector translocating T3SS has been identified in any *Bartonella* species (Fig. 1A). However, our genomic analysis indicated that Bye homologues may have co-evolved with the VirB/VirD4 T4SS and their Bep effectors within lineages 3 and 4 (Fig. 1A). Bye homologues were found in all species of these lineages except for *B. henselae, B. birtlesii* and *B. koehlerae*, suggesting secondary loss by these three species (Fig. 1A). Engel *et al*. (22) proposed that horizontal acquisition of genes encoding the VirB/VirD4 T4SS and at least one Bep by the last common ancestor of lineages 3 and 4 triggered parallel adaptive radiations that gave rise to explosive speciation in these most proliferative lineages. This scenario of evolutionary success due to the acquisition of key innovative traits may be extended to Bye homologues based on their co-appearance with VirB/VirD4 and Beps (Fig. 1A). Independent acquisition of *bye* loci by the last common ancestors of lineages 3 and 4 is supported by the phylogenetic tree of 36 Bye homologues that is monophyletic and largely divided in a lineage 4-specific clade and several lineage 3-specific clades, with evidence for limited horizontal gene transfer between these lineages (Fig. 1B). Moreover, the single *bye* locus of lineage 4 (Fig. 1C) and the multiple *bye* loci of lineage 3 (suppl. Figure S1A-C) display lineage-specific synthenies. Taken together, our evolutionary analysis indicated coevolution of Bye proteins with the VirB/VirD4 T4SS and Bep effectors, suggesting a potential functional link.

**Figure 1.**
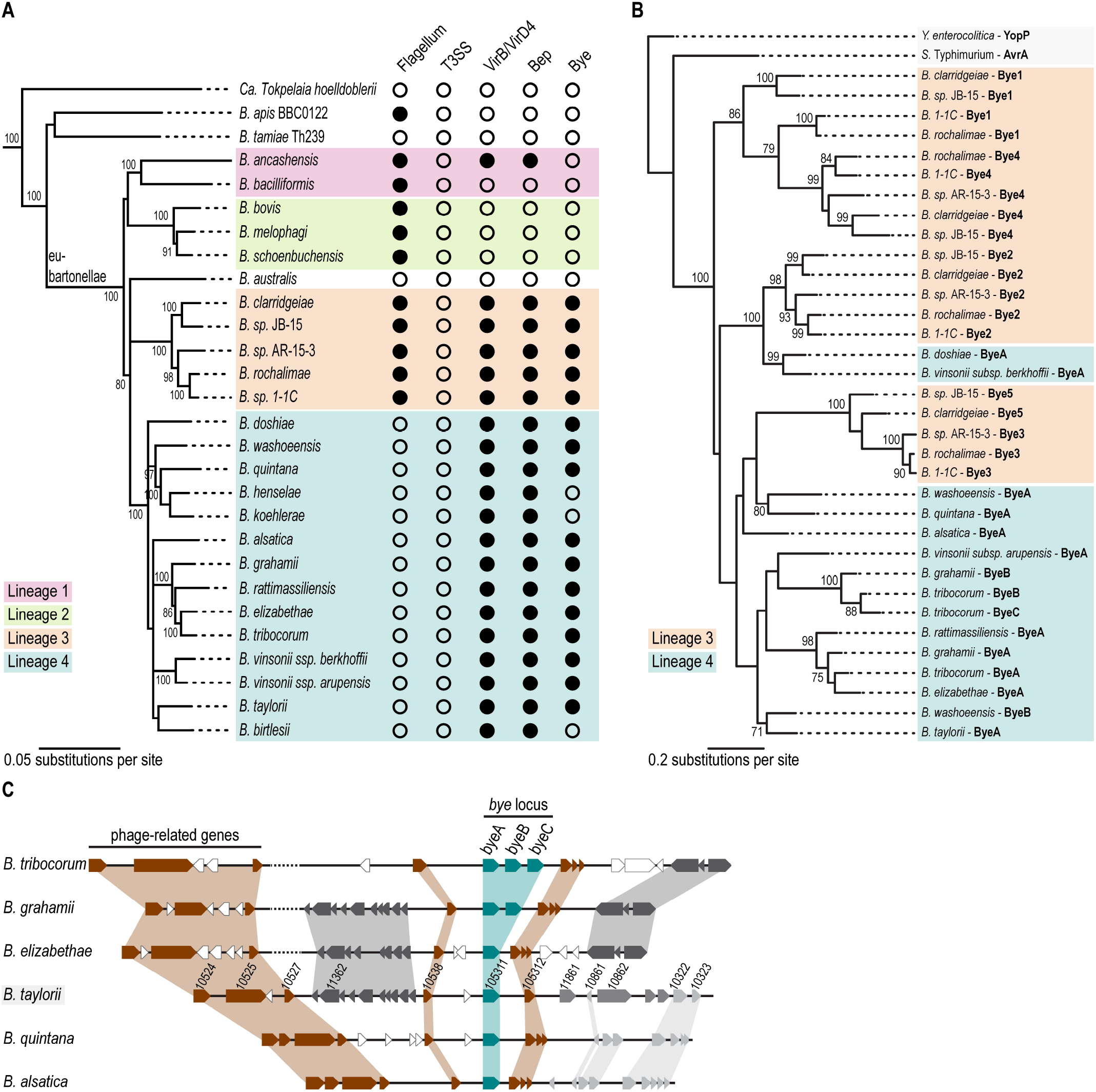
*Bartonella* YopJ-like effector (Bye)-encoding loci have co-evolved with the VirB/VirD4 T4SS. **(A)** Phylogeny of the genus *Bartonella* and distribution of key virulence factors (adapted from Wagner and Dehio, 2019 (17)). The ant-specific species *Candidatus Tokpelaia hoelldoblerii* serves as outgroup taxon. The four *Bartonella* lineages are color coded: lineage 1 in magenta, lineage 2 in green, lineage 3 in orange and lineage 4 in blue. The honeybee symbiont *Bartonella apis* and the pathogenic *Bartonella tamiae* form an ancestral clade, while the position of *Bartonella australis* in the tree is unstable. The phylogenetic tree was inferred based on a concatenated alignment of five core protein sequences (17). The presence and absence of key virulence factors is indicated by full and empty circles, respectively. **(B)** Maximum likelihood tree of the protein sequence alignment of 36 Bye homologs identified in the Bartonellae. The YopJ-family effectors YopP of *Y. enterocolitica* and AvrA of *S.* Typhimurium were included as outgroups to root the tree. Bootstrap values ≥70 of 100 replicates are shown. **(C)** The genomic regions encoding the *bye* loci and flanking regions are compared between six *Bartonella* species of lineage 4 (*B. tribocorum*, *B. grahamii*, *B. elizabethae*, *B. taylorii*, *B. quintana* and *B. alsatica*). Genes are depicted as arrows using the following color code: *bye* homologs (teal), other identified orthologous genes conserved in all compared species (brown), gene orthologues present in some species (grey) and genes without identified orthologues in the other species (white). Gene identifiers are shown for *B. taylorii* (highlighted in grey).

### ByeA impairs MAPK signaling similar to other YopJ family effectors

Next, we aimed to test whether ByeA of *B. taylorii* IBS296 has a similar biological activity as the well-studied YopJ family effectors YopJ/YopP of *Yersinia* or *Salmonella* (AvrA), which are known to potently downregulate innate immune responses. Upon activation of an upstream kinase, the members of the MAPK kinase superfamily and the NF-κB regulatory components IKKα and IKKβ undergo phosphorylation at residues within their activation loop (31, 32). YopJ homologs of *Yersinia* (YopJ/YopP) and *Salmonella* (AvrA) acetylate those residues and thereby block the downstream signaling (5, 11, 33). HeLa cells are a well-established experimental model to study those signaling cascades. Following serum starvation, MAPK and NF-κB components show low levels of phosphorylation, while pro-inflammatory stimuli like LPS of *E. coli* or TNF-α, trigger a transient phosphorylation cascade (5, 34, 35). We transfected HeLa cells with the genes encoding either wild-type ByeA of *B. taylorii* or the predicted catalytically inactive ByeA^C194A^ mutant harboring an amino acid exchange in the conserved catalytic triad. To induce the MAPK and NF-κB pathways we stimulated the cells with TNF-α. We observed phosphorylation of p38, JNK and ERK1/2 MAPK kinases and NF-κB p65 subunit in non-transfected Hela cells and cells expressing catalytically inactive ByeA^C194A^ (Figure 2A and 2B, suppl. Figure S2A and S2B). Phosphorylation of ERK1/2 was not inhibited in ByeA-expressing HeLa cells (suppl. Figure S2A). The phosphorylation of p65 was diminished compared to ByeA^C194A^-expressing cells (suppl. Figure S2B), while phosphorylation of p38 and JNK was virtually abolished in HeLa cells expressing wild-type ByeA (Figure 2A and 2B). The conserved function of ByeA might indicate a contribution to the downregulation of the host’s innate immune responses during *Bartonella* infection. However, given that Bartonellae do not encode effector-translocating T3SSs (18), the mechanism of Bye translocation into host cells remained elusive.

**Figure 2.**
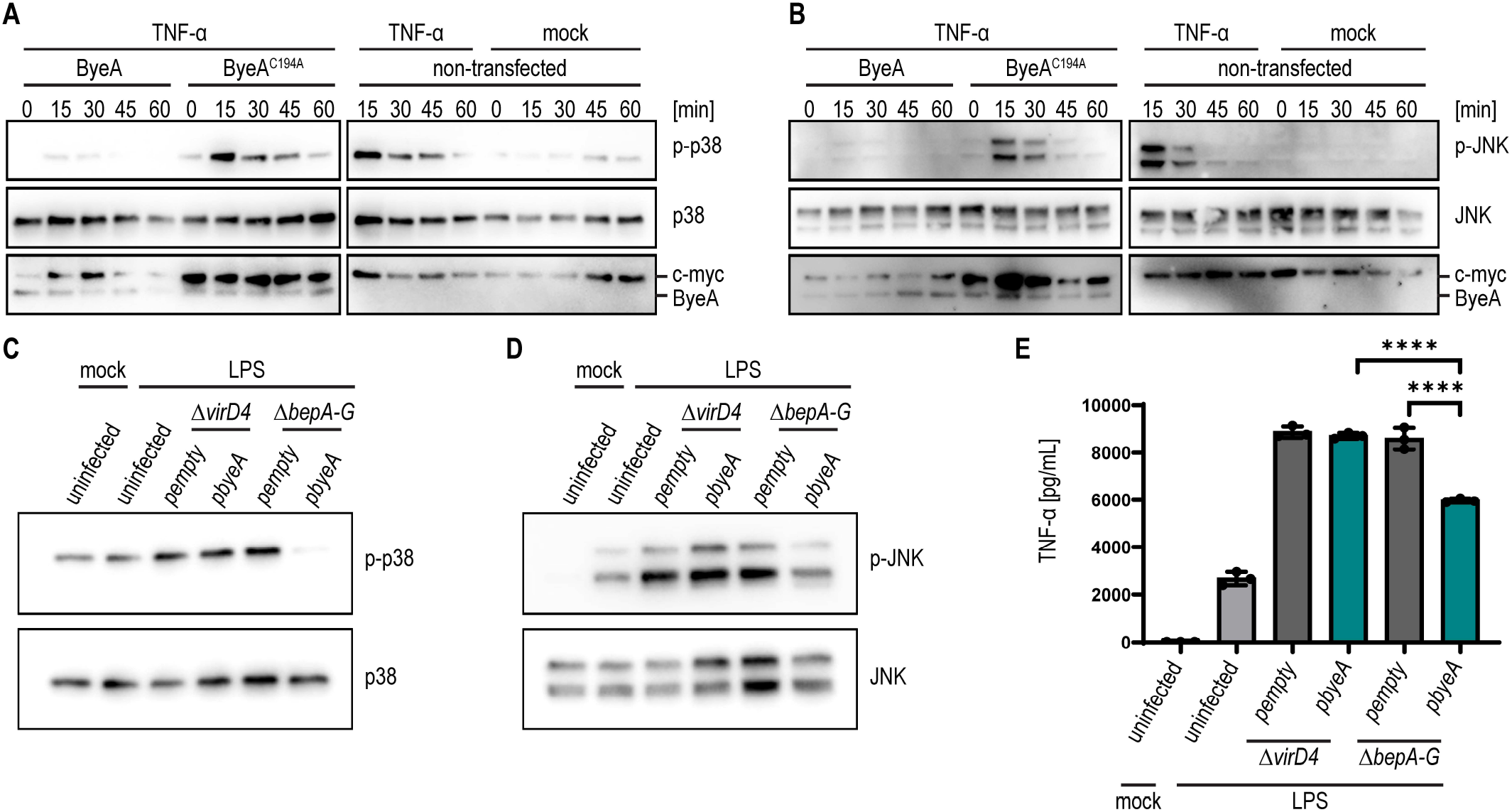
ByeA of *B. taylorii* inhibits TNF-α-induced phosphorylation of p38 and JNK and impairs TNF-α secretion in a VirB/VirD4-dependent manner. **(A, B)** HeLa cells were transiently transfected with expression constructs for ByeA wild-type or the catalytically inactive ByeA^C194A^ mutant proteins carrying an N-terminal c-myc epitope tag. Following 12-16 h of serum starvation, cells were treated with 20 ng/ml human recombinant TNF-α for the indicated times. HeLa cells were harvested, lysed and analyzed by immunoblot using the indicated antibodies. Expression levels of ByeA constructs were monitored with an anti-c-myc tag antibody. **(A)** Phosphorylation of p38 or total p38 levels were monitored using the monoclonal antibodies p-p38 (T180/Y182) or p38, respectively. **(B)** Phosphorylation of JNK or total JNK levels were monitored using the monoclonal antibodies against p-JNK (T183/Y185) or JNK, respectively. **(C-E)** RAW 264.7 macrophages were infected for 5 h at MOI 50 with *B. henselae* wild-type, the Δ*virD4* or Δ*bepA-G* mutants expressing ByeA from a plasmid (p*byeA*) or containing the empty plasmid control (p*empty*). Except for the untreated control, cells were co-stimulated with 100 ng/ml *E. coli* LPS during the last hour of infection. **(C)** Immunoblot analysis using monoclonal antibodies against p-p38 (T180/Y182) and p38. **(D)** Immunoblot analysis using monoclonal antibodies against p-JNK (T183/Y185) and JNK. **(E)** Secreted TNF-α was quantified by ELISA. All data show representative results from three independent experiments. **(E)** shows the mean ±SD of technical triplicates. Data was analyzed using one-way ANOVA with multiple comparisons (Tukey’s multiple comparison test), **** p < 0.0001.

### ByeA downregulates TNF-**α** secretion in a VirB/VirD4-dependent manner

*B. henselae* and *B. taylorii* were shown to translocate *Bartonella* effector protein D (BepD) via the VirB/VirD4 T4SS resulting in downregulation of pro-inflammatory cytokine (*e.g.,* TNF-α) secretion (36). However, mutant analysis indicated that *B. taylorii* must translocate at least one additional anti-inflammatory effector via its VirB/VirD4 T4SS that is different from the known Bep effectors (BepA-I) (37). As ByeA of *B. taylorii* inhibits p38 and JNK signaling, which might downregulate pro-inflammatory cytokine secretion as shown for other YopJ-family effectors, ByeA represents a candidate for this elusive immune-modulatory effector. Therefore, we tested if ByeA is translocated via the VirB/VirD4 T4SS.

*B. henselae* is the best-characterized *Bartonella* species for functional studies on the VirB/VirD4 T4SS and Bep translocation into eukaryotic host cells (36–38) and does not encode a Bye homologue (Fig. 1A). To test for inhibition of the p38 and JNK signaling pathway in dependence of the VirB/VirD4 T4SS, we investigated their phosphorylation in RAW 264.7 macrophages infected with *B. henselae* mutant strains ectopically expressing ByeA of *B. taylorii*. To trigger MAPK pathways, we stimulated the cells with LPS during the last hour of infection. We found that a *B. henselae* strain deleted for all endogenous Beps (Δ*bepA-G*, effector-less) that ectopically expresses ByeA from a plasmid (p*byeA*) decreases the phosphorylation of p38 and JNK at 5 hpi (Figure 2C and 2D) and 24 hpi (suppl. Figure S2C and S2D). The translocation-deficient mutant Δ*virD4* expressing ByeA from a plasmid (p*byeA*) or the effector-less mutant Δ*bepA-G* carrying the empty expression plasmid (p*empty*) did not affect phosphorylation. The secretion of pro-inflammatory TNF-α was significantly reduced after infection with *B. henselae* Δ*bepA-G* p*byeA* but not Δ*virD4* p*byeA* at 5 hpi (Figure 2E) and 24 hpi (suppl. Figure S2E). Mutant analysis in *B. taylorii* confirmed a role for ByeA in downregulating pro-inflammatory TNF-α secretion (suppl. Figure S3). Deletion of *byeA* resulted in increased TNF-α secretion, while ectopic expression of ByeA from a plasmid in different deletion backgrounds reduced TNF-α secretion close to the level of the wild-type (suppl. Figure S3C). Taken together, our data revealed that ByeA of *B. taylorii* suppresses innate immune responses in a VirB/VirD4 T4SS-dependent manner.

### ByeA translocation via the VirB/VirD4 T4SS of *Bartonella* depends on conserved regulatory sequences flanking the acetyltransferase domain

We next characterized the T4SS-specific translocation signal of *B. taylorii* ByeA using the recently established split NanoLuc luciferase-based translocation assay (37, 39). In short, the large fragment of NanoLuc (LgBiT) is stably expressed in RAW 264.7 macrophages, while the small fragment (HiBiT) is expressed in isogenic *B. henselae* strains as a FLAG-tagged N-terminal fusion to sequences of ByeA. While both fragments have no luciferase activity on their own, the VirB/VirD4-dependent translocation of a HiBiT fusion protein into the RAW LgBiT macrophages leads to the reconstitution of a functional luciferase that upon substrate administration generates a luminescent signal (Figure 3A).

**Figure 3.**
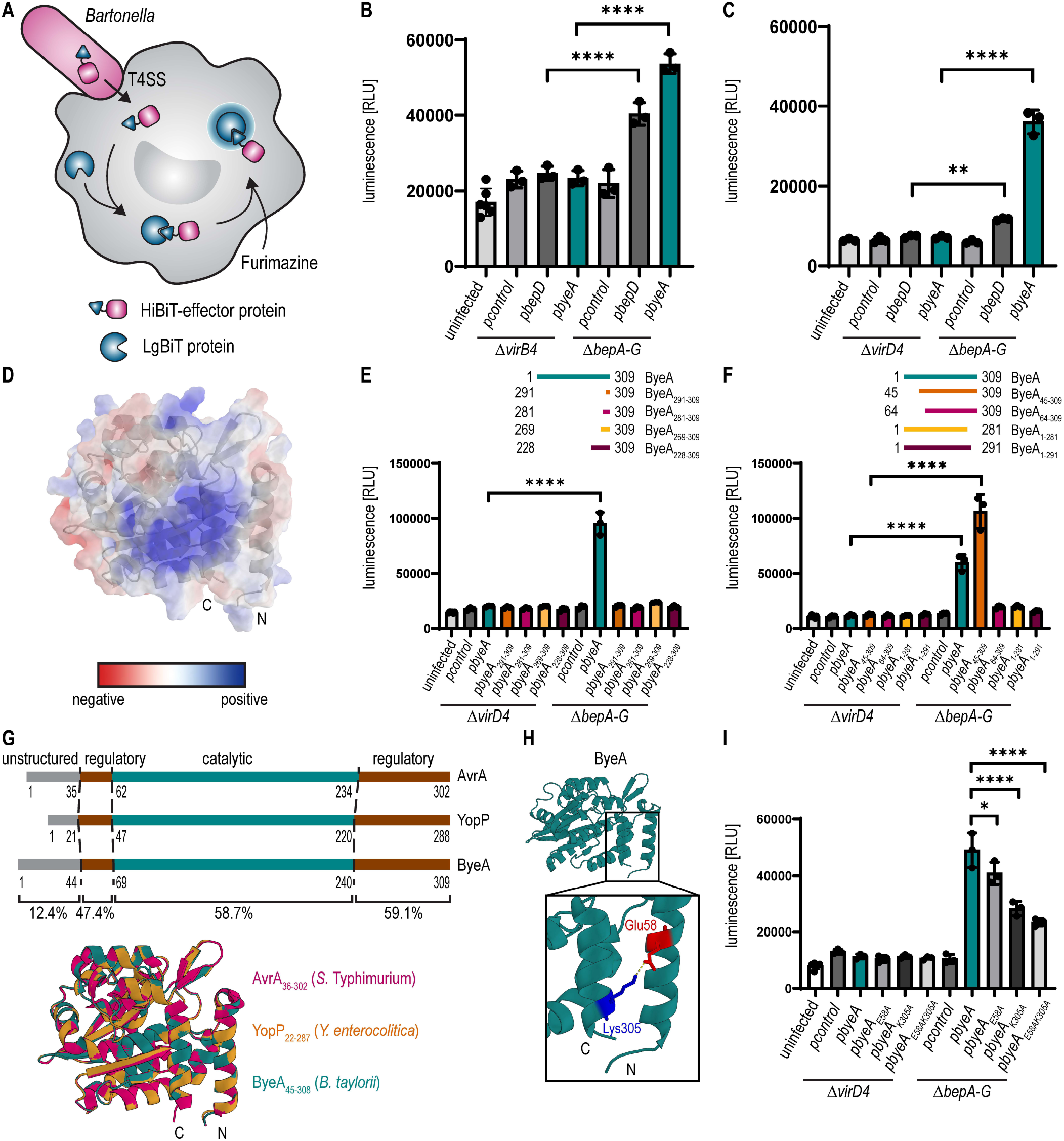
The C-terminus and an N-proximal helix are important for the translocation of ByeA. **(A)** Schematic representation of the split NanoLuc luciferase-based translocation assay. **(B, C)** *B. henselae* Δ*virB4*, Δ*virD4* or Δ*bepA-G* mutants containing the indicated plasmid were incubated for 24 h in M199 + 10% FCS at 35°C and 5% CO_2_. To induce protein expression, the cultures were supplemented with 100 μM IPTG for at least 60 min prior to infection. For simplicity, the *hibit-flag* sequence, which is encoded by p*control* and N-terminally fused to *bepD* and *byeA*, is not indicated in the plasmid names. RAW LgBiT macrophages were infected with *B. henselae* strains for 24 h followed by supplementation with the NanoLuc substrate. **(B)** Cells infected with *B. henselae* Δ*virB4* or Δ*bepA-G* mutants containing the indicated plasmids. **(C)** Cells infected with *B. henselae* Δ*virD4* or Δ*bepA-G* mutants containing the indicated plasmids. **(D)** Electrostatic potential of charged amino acids shown for ByeA using the “APBS Tool2.1” plugin in PyMOL (version 2.3.4). **(E, F, I)** For simplicity, the *hibit-flag* sequence, which is encoded by p*control* and N-terminally fused to all *byeA* sequences, is not indicated in the plasmid names. **(E)** RAW LgBiT macrophages were infected with *B. henselae* Δ*virD4* or Δ*bepA-G* mutant backgrounds expressing different C-terminal regions of ByeA as depicted in the legend. **(F)** RAW LgBiT macrophages were infected with *B. henselae* Δ*virD4* or Δ*bepA-G* mutant backgrounds expressing different ByeA truncations as depicted in the legend. **(G)** Domain architecture of ByeA, YopP and AvrA with the unstructured N-terminus (grey), regulatory domains (brown) and catalytic core (teal). Pairwise identity of the three homologs is given in percentage for the different domains. The structures of ByeA (shown in teal) and YopP (shown in orange) were predicted using PyMOL version 2.3.4 with the structure of AvrA^ΔL140^ (PDB: 6BE0; shown in pink) as reference. The first 35 amino acids of AvrA have not been solved in the crystal structure and are depicted as unstructured N-terminus. The boundaries of the regulatory and catalytic domains depicted for YopP and ByeA were identified using the published domains of AvrA as reference. **(H)** Structure of ByeA was modeled using PyMOL (version 2.3.4) with focus on the C-terminal and N-terminal helices. Positively charged amino acid (Lys305) is depicted in blue, negatively charged amino acid (Glu58) shown in red. Predicted salt bridge is shown in yellow. **(I)** RAW LgBiT macrophages were infected with *B. henselae* Δ*virD4* or Δ*bepA-G* mutants expressing ByeA or different single amino acid mutants of ByeA (ByeA_E58A_, ByeA_K305A_, ByeA_E58AK305A_). All data show representative results from three independent experiments. **(B, C, E, F, I)** show the mean ±SD of technical triplicates. Data was analyzed using one-way ANOVA with multiple comparisons (Tukey’s multiple comparison test), * p < 0.05, ** p < 0.01, *** p < 0.001, **** p < 0.0001.

To test for functionality of a ByeA full-length fusion in this assay we introduced plasmids encoding HiBiT-FLAG (p*control*, negative control), HiBiT-FLAG-BepD (p*bepD*, *bepD* sequence of *B. henselae*, positive control (37)) or HiBiT-FLAG-ByeA (p*byeA, byeA* sequence of *B. taylorii*) into the *B. henselae* effector-less Δ*bepA-G* background or the T4SS-deficient Δ*virB4* background.

Expression of the HiBiT-FLAG-effector fusions was confirmed by anti-FLAG immunoblot of bacterial lysates (suppl. Figure S4A). The capacity to complement LgBiT to a functional luciferase was confirmed by supplementing bacterial lysates with purified LgBiT and substrate (suppl. Figure S4B). RAW LgBiT macrophages were infected at a multiplicity of infection (MOI) of 50 and the bioluminescent signal was analyzed at 24 hours post infection (hpi). Compared to infections with strains of the Δ*virB4* mutant background, cells infected with Δ*bepA-G* expressing either HiBiT-FLAG-BepD or HiBiT-FLAG-ByeA emitted a significantly increased luminescence signal (Figure 3B). Infection of RAW LgBiT macrophages with *B. henselae* strains lacking the T4CP (Δ*virD4*), did not generate a luminescent signal (Figure 3C). Expression of HiBiT-FLAG-effector fusions and the capacity to complement LgBiT in lysates was confirmed for all strains (suppl. Figure S4B and S4C). In conclusion, translocation of ByeA is dependent on VirB4 and the T4CP VirD4.

Based on this functional translocation assay we characterized the nature of the T4SS-specific translocation signal of ByeA. In most T4SS effectors, the translocation signal localizes to the C-terminus and contains an accumulation of positively charged or hydrophobic amino acids (1, 26, 40). Analyzing a ByeA molecular model (Figure 3D) based on the solved structure of AvrA of *S.* Typhimurium (9), we noted an accumulation of positively charged amino acids (aa) in the C-terminus. In the Beps and related T4SS effectors this C-terminal tail sequence is extended N-terminally by the BID domain (27, 41, 42). To examine if a C-terminal fragment of ByeA may be sufficient to mediate translocation we generated HiBiT-FLAG-fusions fused to C-terminal ByeA fragments of either 18 aa, 28 aa, 40 aa or 81 aa. Most likely due to their low molecular mass (below 14 kDa) these fusion proteins were not detectable by immunoblot analyses. However, in bacterial lysates supplemented with LgBiT all constructs displayed luciferase activity (suppl. Figure S5A), demonstrating their functional expression. Infection of RAW LgBiT macrophages with *B. henselae* Δ*bepA-G* expressing the indicated fusion proteins did not increase the luminescent signal compared to isogenic strains in the translocation-deficient Δ*virD4* mutant (Figure 3E), indicating that the C-terminus of ByeA is insufficient to mediate translocation. The T4SS signal of ByeA thus seems to be fundamentally different from that of other known T4SS effectors.

As we failed to define a region of ByeA that is sufficient for mediating VirB/VirD4-dependent translocation, we aimed to identify regions of the effector that are required for translocation. To this end we constructed C-terminal HiBiT-FLAG fusions to an ordered series of N- or C-terminal truncations of ByeA and expressed them in either the effector-less Δ*bepA-G* background or the translocation-deficient Δ*virD4* background (Figure 3F). We confirmed expression of the fusions by immunoblot analysis and demonstrated in bacterial lysates their capacity to complement LgBiT to a functional luciferase (suppl. Figure S5B and S5C). Compared to infections with bacteria expressing full-length ByeA (p*byeA*), deletion of the first 44 aa (p*byeA_45-309_*) even increased the luminescent signal, while deletion of the first 63 aa (p*byeA_64-309_*) resulted only in background signal (Figure 3F). The N-terminal 44 aa are thus dispensable for translocation, or even inhibit it. The corresponding N-terminal regions are highly variable in size and amino acid composition among the various ByeA homologues of lineage 3 and 4 and unrelated to corresponding regions of YopP and AvrA (suppl. Figure S6A and S7A). In contrast, the region from aa 45 to 63 is strictly required for translocation. Infection with Δ*bepA-G* strains expressing C-terminally truncated proteins (p*byeA_1-291_* and p*byeA_1-281_*) showed only background luminescence (Figure 3F), indicating that sequences within the C-terminal 18 aa (positions 282-309) are essential for translocation via the VirB/VirD4 T4SS.

Importantly, the regions identified as essential for VirB/VirD4-dependent translocation proximal to the N-terminus (positions 45-63) or at the C-terminus (position 282-309) are fully comprised within highly conserved regulatory regions flanking the central acetyltransferase region either on the N-terminal (positions 45-69, 47.4% identity with YopP and AvrA) or C-terminal side (positions 241-309, 59.1% identity with YopP and AvrA) (Figure 3G and suppl. Figure S6A). These regulatory regions comprise most of the residues involved in binding of the cofactors IP6 and acetyl-CoA, previously identified for AvrA and HopZ1a of *Pseudomonas syringe* (8, 9) (suppl. Figure S6A and S7A-C). Based on homology modeling of ByeA, YopP and AvrA the protein structures appear highly conserved among the regulatory regions and central acetyltransferase domain, while the N-termini are unstructured and could not be resolved in the crystal structure of AvrA. The regulatory regions in the N- and C-terminus form α-helices and in the folded protein these helices are aligned in an anti-parallel orientation (Figure 3G). The homology modeling further predicted that a negatively charged glutamate in the helix proximal to the N-terminus interacts with a positively charged lysine in the C-terminal helix (ByeA: E58 and K305; YopP: E36 and K283; AvrA: E50 and K297) (Figure 3H, suppl. Figure S6B). To investigate whether this putative salt bridge between the two anti-parallel helices may be critical for ByeA translocation, we created point mutations that convert the charged amino acids to an alanine. We introduced respective plasmids into the effector-less Δ*bepA-G* background or the translocation-deficient Δ*virD4* background (Figure 3I). We confirmed expression of the fusions by immunoblot analysis and demonstrated in bacterial lysates their capacity to complement LgBiT to a functional luciferase (suppl. Figure S6C and S6D). Compared to infection of RAW LgBiT macrophages with *B. henselae* Δ*bepA-G* expressing ByeA wild-type (p*byeA*), Δ*bepA-G* expressing HiBiT-FLAG-ByeA_E58A_ (p*byeA_E58A_*) resulted in lower luminescent signals. The bioluminescence was further decreased for Δ*bepA-G* expressing HiBiT-FLAG-ByeA_K305A_ (p*byeA_K305A_*) and dropped even further when the double-mutant, HiBiT-FLAG-ByeA_E58AK305_ (p*byeA_E58AK305A_*), was expressed (Figure 3I). These mutant analysis data imply a critical role for both amino acids E58 and K305 for ByeA translocation, which may go beyond a potential structural role in mediating ionic interaction. Taken together, both helices localizing proximal to the N-terminus and at the C-terminus are essential to mediate translocation through the VirB/VirD4 T4SS.

### T3SS-translocated YopJ-family effectors are also translocated via the T4SS of *Bartonella*, while ByeA is not translocated via the T3SS of *Yersinia*

The proximal to N-terminus and the C-terminal sequences of ByeA required for T4SS-dependent translocation in *Bartonella* are located in regulatory sequences generally conserved among YopJ-family effectors. Therefore, we wondered whether the T3SS effectors YopP and AvrA may also carry a functional T4SS-translocation signal as demonstrated for ByeA. We used the split NanoLuc translocation assay to test whether YopP and AvrA are translocated through the VirB/VirD4 T4SS of *Bartonella*. To this end we expressed N-terminal HiBiT-FLAG-fusions to either ByeA, YopP or AvrA in *B. henselae* in the effector-less Δ*bepA-G* or the translocation-deficient Δ*virD4* mutant. Protein expression or functional complementation with LgBiT to a functional luciferase were tested by immunoblot or luciferase assay, respectively (suppl. Figure S8A and S8B). MOI screens revealed that infection of RAW LgBiT macrophages at MOI ≥ 50 with Δ*bepA-G* containing p*hibit-flag-byeA* (p*byeA*), p*hibit-flag-yopP* (p*yopP*) or p*hibit-flag-avrA* (p*avrA*) significantly increased the luminescent signal compared to the isogenic strains in the Δ*virD4* mutant background (Figure 4A-C). These data demonstrate that YopP and AvrA are translocated via the VirB/VirD4 T4SS of *Bartonella,* albeit with lower efficiency than ByeA.

**Figure 4.**
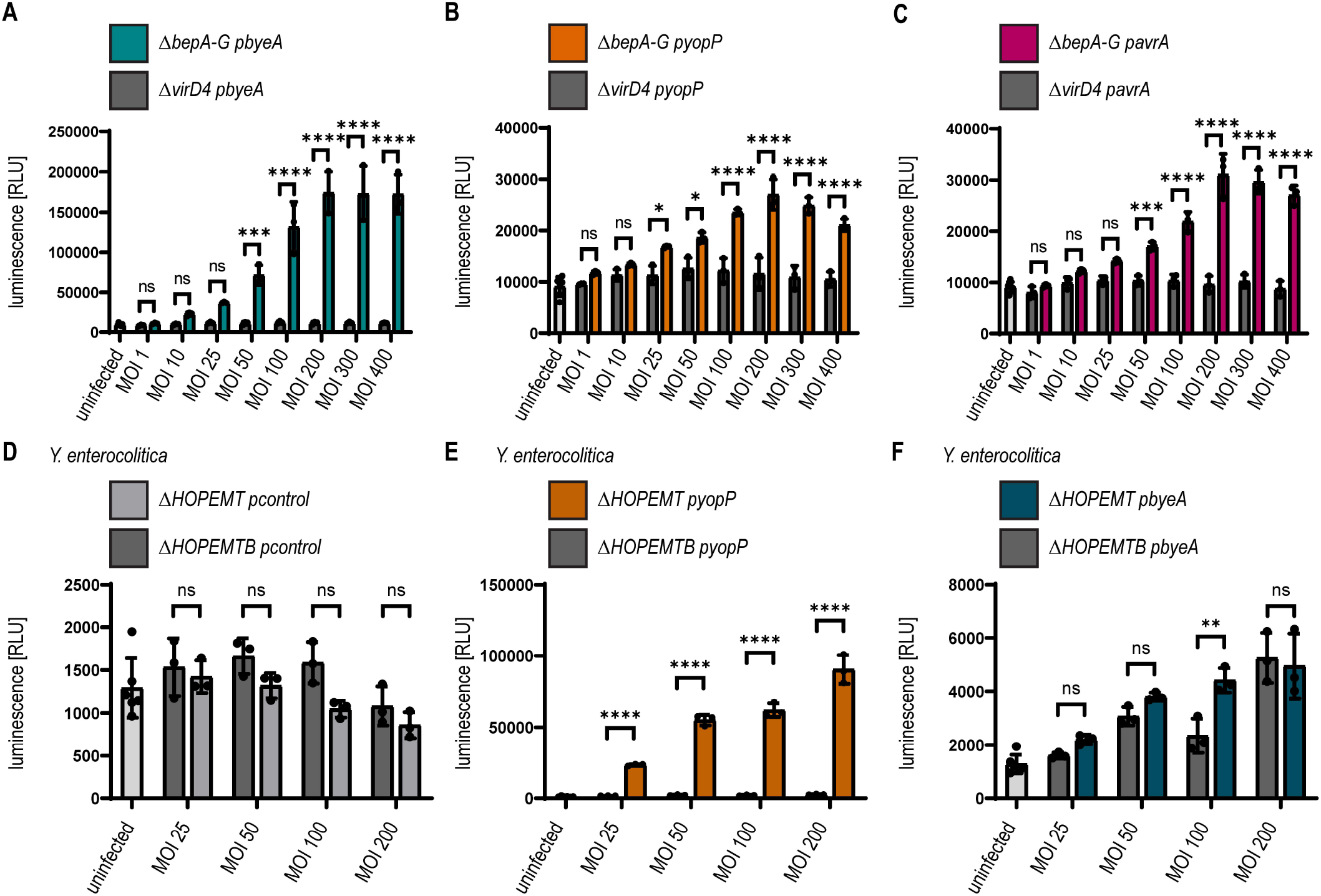
The YopJ-family T3SS effectors YopP and AvrA are secreted via the T4SS of *B. henselae*, while the homologous ByeA is not translocated via the T3SS of *Y. enterocolitica*. **(A-C)** *B. henselae* Δ*virD4* or Δ*bepA-G* mutants were incubated for 24 h in M199 + 10% FCS at 35°C and 5% CO_2_. To induce protein expression, the cultures were supplemented with 100 μM IPTG for at least 1 h prior to infection. For simplicity, the *hibit-flag* sequence, which is N-terminally fused to all depicted effectors in *Bartonella*, is not indicated in the plasmid names. RAW LgBiT macrophages were infected using the depicted MOIs with (A) *B. henselae* Δ*virD4* or Δ*bepA-G*, both expressing HiBiT-FLAG-ByeA (p*byeA*), (B) *B. henselae* Δ*virD4* or Δ*bepA-G* expressing HiBiT-FLAG-YopP (p*yopP*) or (C) *B. henselae* Δ*virD4* or Δ*bepA-G* expressing HiBiT-FLAG-AvrA (p*avrA*). (D-F) *Y. enterocolitica* effector-less mutant Δ*HOPEMT* and the translocation-deficient effector-less mutant Δ*HOPEMTB* were supplemented with 0.2% arabinose to induce protein expression. For simplicity, the *flag-hibit* sequence, which is C-terminally fused to all depicted effectors in *Yersinia*, is not indicated in the plasmid names. RAW LgBiT macrophages were infected using the depicted MOIs for 2 h with (D) *Y. enterocolitica* Δ*HOPEMT* or Δ*HOPEMTB* expressing FLAG-HiBiT (p*control*), (E) *Y. enterocolitica* Δ*HOPEMT* or Δ*HOPEMTB* expressing YopP-FLAG-HiBiT (p*yopP*) or (F) *Y. enterocolitica* Δ*HOPEMT* or Δ*HOPEMTB* expressing ByeA-FLAG-HiBiT (p*byeA*) **(A-F)** show the mean ±SD of technical triplicates. The luminescence of the complemented split NanoLuc was measured. Data was analyzed using one-way ANOVA with multiple comparisons (Tukey’s multiple comparison test), ns = not significant, * p < 0.05, ** p < 0.01, *** p < 0.001 **** p < 0.0001.

We then used the split NanoLuc translocation assay reciprocally to test if ByeA may also be translocated by the T3SS of *Y. enterocolitica*. As the T3SS secretion signal is located N-terminally, we fused the FLAG-tagged HiBiT fragment C-terminally to either YopP, serving as positive control, or ByeA. *Yersinia* lacking the effectors YopH, YopO, YopP, YopE, YopM and YopT (Δ*HOPEMT*) and the isogenic translocation-deficient bacteria lacking the translocator YopB (Δ*HOPEMTB*) (43, 44) were transformed with a plasmid encoding *flag*-tagged *hibit* (p*control*), p*byeA-flag-hibit* (p*byeA*) or p*yopP-flag-hibit* (p*yopP*). Expression of the FLAG-HiBiT fusion proteins or complementation with LgBiT to a functional luciferase were examined, respectively (suppl. Figure S8C and S8D). We did not detect any significant differences in the luminescent signal following the infection of RAW LgBiT macrophages with Δ*HOPEMT* containing p*flag-hibit* compared to the isogenic translocation-deficient mutant Δ*HOPEMTB* (Fig. 4D). Infection of RAW LgBiT macrophages with Δ*HOPEMT* containing p*yopP-flag-hibit* significantly increased the luminescent signal over the isogenic translocation-deficient strain Δ*HOPEMTB*, demonstrating functionality of the split NanoLuc translocation assay for the endogenous T3SS substrate YopP (Figure 4E). In contrast, cell infection with Δ*HOPEMT* containing p*byeA-flag-hibit* did not consistently display elevated luminescence over the Δ*HOPEMTB* strain, indicating that ByeA is unlikely translocated via the T3SS of *Y. enterocolitica* (Figure 4E, Figure S8G and J). Nonetheless, we observed an increase in luminescence after infection with higher MOIs for both strains, Δ*HOPEMT* and Δ*HOPEMTB,* harboring p*byeA-flag-hibit*. This increase in luminescence, which is independent of the T3SS, may be attributed to partial cell lysis.

Taken together, our data show that the genuine T3SS effectors YopP and AvrA are also translocated via the T4SS of *Bartonella*, while the genuine T4SS effector ByeA is unlikely translocated by the T3SS of *Yersinia*.

## Discussion

Many bacterial pathogens use dedicated secretion systems to translocate effector proteins into host cells in order to modify host cell functions to their benefit (45). Most translocated effectors contain a secretion system-specific translocation signal and typically one or several functional domains that interfere with specific host cell processes (3). Among the many functional domains described in bacterial effectors, some were shown to be derived from pre-evolved domains of prokaryotic or eukaryotic origin (3, 45), while some others evolved apparently *de novo* as part of translocated effectors (46). In some instances, homologous domains have been found to be shared by effector proteins translocated via different bacterial secretion systems. However, their evolutionary history is often unclear. In a parallel evolutionary scenario, these effectors evolved independently from each other by fusion of a pre-evolved functional domain to different translocation signals. Examples are the FIC domains found in several T3SS and T4SS effector families that derived from diversified members of the pervasive family of FicT/A toxin-antitoxin systems present in many bacteria (41, 47). In a different scenario of divergent evolution, an effector domain evolved initially in association with a given translocation signal then becomes secondarily associated with a translocation signal specific for another secretion system. Such effector switches for secretion system specificity could result from horizontal gene transfer and illegitimate recombination, but evidence is scarce as the evolutionary history can only be inferred if the original translocation signal does not deteriorate upon the acquisition of a different translocation signal. Studying the pervasive YopJ family of effector proteins, we have described here such a switch of secretion system specificity.

YopJ family effector proteins belong to the best studied host-targeted bacterial effectors (4). Up to now they were recognized as a widely distributed family of closely related T3SS-translocated effectors, containing a poorly conserved N-terminal T3SS translocation signal and a conserved central acetyltransferase domain. The enzymatic activity of the acetyltransferase domain is controlled by flanking regulatory sequences. In particular, the binding of IP6 to these sequences induces a conformational change that is required for binding of acetyl-CoA that then serves as acetyl donor in the acetylation reaction (8). Studying translocation via the VirB/VirD4 T4SS of *B. henselae* we found that these conserved sequences, which regulate the acetyltransferase activity, coincide with a co-evolved T4SS translocation signal. Identified initially in the novel T4SS effector ByeA of *B. taylorii*, the T4SS translocation signal is also functionally conserved in the well characterized T3SS effectors YopP of *Y. enterocolitica* and AvrA of *Salmonella* Typhimurium. Given that (i) ByeA of *B. taylorii* is apparently not translocated via the T3SS of *Y. enterocolitica* and that (ii) ByeA homologues are limited to *Bartonella* lineages that encode a VirB/VirD4 T4SS, it is tempting to speculate that ByeA homologs may represent an ancestral state of genuine T4SS effectors. The fusion of an N-terminal T3SS signal may then have given rise to the evolution of the large group of T3SS effectors dramatically expanding the YopJ family, with the T4SS translocation signal remaining functional conserved due to their coincidence with co-evolved regulatory sequences for the acetyltransferase activity. While we consider this evolutionary scenario to be the most plausible, it is important to acknowledge the alternative possibility that a YopJ family effector, which originally functioned as a genuine T3SS effector, may have been acquired by *Bartonella* through horizontal gene transfer. In this alternative scenario, the *Bartonella* T4CP VirD4 would need to have been pre-adapted for effective interaction with the YopJ family effector through the conserved regulatory sequences within the acetyltransferase domain. Several studies have shown that T4CPs can recognize substrates using various signals, e.g. the internal TSA and TSB domains of conjugative relaxases (48, 49), some positively charged amino acids in the C-terminus of *A. tumefaciens* effector proteins (26) or conserved C-proximal motifs of *Xanthomonas citri* protein toxins (40, 50). Noteworthy, some bacteria encoding *yopJ* homologs harbor T3SSs and T4SSs, like *Xanthomonas* or certain *Salmonella* serovars (51, 52). In these species YopJ homologs are translocated by the appropriate T3SS (53–57), however studies showing a potential interaction with the present T4SS were not conducted.

Further to translocating the novel ByeA effector, the *Bartonella* VirB/VirD4 T4SS was previously reported to translocate cocktails of *Bartonella* effector proteins (Beps) (21). While diverse in structure and function, all Beps contain at their C-termini a bi-partite translocation signal that is composed of the conserved BID domain flanked by a positively charged tail sequence (21, 27). In contrast, the T4SS signal of YopJ family effectors as investigated in detail for ByeA of *B. taylorii* is more complex. It comprises an alpha-helix locating to the very C-terminus as well as another one locating to a position proximal to the N-terminus, which align with each other to form an anti-parallel two-helix bundle. Modeling of the highly conserved structural folds of ByeA, YopP and AvrA indicated that this translocation signal is part of one compact fold encompassing the entire protein and has as such limited resemblance to the structure of the BID domain (27). While the sequence similarity of several BID domains is rather low, the surface charge distribution appears consistent with two highly positively charged areas and a negative patch in the center (27, 42). In accordance, we identified a cluster of positively charged amino acids in the C-terminus of ByeA. The mainly positively charged surface suggests that T4SS effectors might interact with a negatively charged interaction partner. The all-alpha domain (AAD) of the VirD4 coupling protein locates at the cytosolic pole of the T4SS channel and displays a negatively charged surface distribution (58, 59), which might be crucial for the recognition of positively charged secretion signals of T4SS effectors (27, 42). Future structure-function studies should address how VirD4 recognizes these distinctive translocation signals at the molecular level during the initial step in effector translocation via the VirB4/VirD4 T4SS.

In summary, our functional and evolutionary studies on YopJ family proteins establish an example of a bacterial effector switching secretion system-specificity, which might represent a more frequent process than presently appreciated and adds new aspects to our knowledge of the nature of the T4SS translocation signal.

## Materials and Methods

More detailed information can be found in supplementary material.

### Bacterial strains and growth conditions

*Escherichia coli*, *Bartonella* strains and *Yersinia* strains were cultured as previously described (37, 60). See supplementary material for more detailed information. Bacterial strains used in this study are listed in supplementary table S1.

### Construction of strains and plasmids

The sequences of all oligonucleotide primers used in this study are listed in supplementary table S2. A detailed description for the construction of each plasmid is presented in supplementary table S3.

### Cell lines and culture conditions

The murine macrophage cell line RAW 264.7 (ATCC TIB-71, (61)) and RAW LgBiT macrophages (37) were cultured at 37°C and 5% CO_2_ in DMEM Glutamax supplemented with 10% heat-inactivated FCS. RAW LgBiT cells were treated with 2 ng/ml puromycin to select for stably transduced cells. HeLa cells (ATCC® CCL-2™, (62)) were cultured in DMEM Glutamax supplemented with 10% heat-inactivated FCS at 37°C and 5% CO_2_.

### Cellular infection

See supplementary materials for a detailed description of infection of RAW 264.7 and RAW LgBiT macrophages with either *Bartonella* or *Yersinia* strains.

### Transient transfection of HeLa cells and determination of ByeA targets

HeLa cells were transfected with plasmids encoding ByeA or ByeA^C197A^ according to the FuGENE transfection protocol (FuGENE® HD Transfection Reagent, Promega Cat. E2311) as advised by the manufacturer. Activation of MAPK and NF-κB signaling pathways was triggered using human recombinant TNF-α (BioLegend, Cat. 570102). Detailed information concerning experimental procedure is given in supplementary material.

### NanoLuc-based effector translocation assay

To assess whether the HiBiT-FLAG fragment or effectors fused to HiBiT-FLAG can complement LgBiT to a functional luciferase, we used the Nano-Glo HiBiT lytic detection system (Promega, Catv N3030). To determine the linear range of the luminescence signal we performed a dilution series with different numbers of bacteria. The Nano-Glo live cell reagent (Promega) was used as advised by manufacturer to determine effector translocation into RAW LgBiT macrophages. A more detailed description of experimental procedure and instrument settings can be found in supplementary material.

### SDS-Page and Immunoblot analysis

SDS-PAGE and immunoblotting were performed as described (21). See supplementary materials for a detailed experimental procedure and listed antibodies to detect proteins of interest.

### Quantification of secreted TNF-**α**

TNF-α concentration in cell culture supernatants was determined using Ready-SET-Go! ELISA kits (ThermoFisher, Cat. 88-7324-88) according to the manufacturer’s instruction. For further details, refer to supplementary material.

### Bioinformatic analysis

A detailed description of bioinformatics analysis can be found in supplementary material. The database from annotated *Bartonella* species can be found in supplementary table S5.

### Statistical analysis

Graphs were generated with GraphPad Prism 9. Statistical analyses were performed using one-way ANOVA with multiple comparisons (Tukey’s multiple comparison test). For the graphs presented in the figures, significance was denoted as non-significant (ns) (p > 0.05); * p < 0.05; ** p < 0.01; *** p < 0.001; P**** < 0.0001. Number of independent biological replicates is indicated as n in the figure legends.

## Acknowledgments

We thank T3 Pharmaceuticals, Allschwil, Switzerland, for providing the *Yersinia* strains and supporting us during the related infections assays. We especially want to thank Lena Siewert, Jaroslaw Sedzicki for technical support, and Lena Siewert, Jaroslaw Sedzicki and Isabel Sorg for helpful comments and critical reading of the manuscript. Portions of the paper were developed from the thesis of K. F.

