## Supplemental Figures and Tables for "Translocation of YopJ family effector proteins through the VirB/VirD4 T4SS of *Bartonella*"

### **Supplementary Information for**
Translocation of YopJ family effector proteins through the VirB/VirD4 T4SS of *Bartonella*

Katja Fromm, Monica Ortelli, Alexandra Boegli, Christoph Dehio

Christoph Dehio


**This PDF file includes:**

- Supplementary Material & Methods
- Figures S1 to S8
- Tables S1 to S4
- SI References

#### Supplementary Material & Methods

##### Bacterial strains and growth conditions

Bacterial strains used in this study are listed in supplementary Table S1. *E. coli* strains were cultivated on solid agar plates (LA) or in lysogeny broth (LB) supplemented with the appropriate antibiotics at 37°C.

*Bartonella* strains were grown on Columbia blood agar (CBA) or tryptic soy agar (TSA) plates supplemented with 5% defibrinated sheep blood (Thermo Scientific Oxoid, Cat. SR0051) and the appropriate antibiotics at 35°C and 5% $CO_2$. *Bartonella* strains were stored as frozen stocks at -80°C and inoculated on CBA or TSA plates for 3 days. Subsequently, the bacteria were expanded on fresh plates for 2 days. Prior to infection, *Bartonella* strains were inoculated at an optical density ($OD_{600nm}$) of 0.5 in M199 medium supplemented with 10% heat-inactivated fetal calf serum (FCS) and incubated for 24 h at 35°C (*B. henselae*) or 28°C (*B. taylorii*) and 5% $CO_2$.

*Yersinia* strains were cultured in brain heart infusion (BHI) medium or on LA plates at room temperature (RT) supplemented with appropriate antibiotics. Prior to infection, bacterial overnight cultures were diluted to an $OD_{600nm}$ of 0.2 and shifted into a 37°C water bath shaker for 2 h to induce the assembly of the T3SS (1).

Antibiotics or supplements were used in the following concentrations: streptomycin at 100 µg/ml, ampicillin at 200 ng/ml, kanamycin at 30 µg/ml, isopropyl-β-D-thiogalactoside (IPTG) at 100 µM, nalidixic acid (Nal) at 35 µg/ml, arsenite at 400 µM, 0.2% arabinose and diaminopimelic acid (DAP) at 1 mM.

##### Construction of strains and plasmids

DNA manipulations were performed according to standard techniques and inserts were sequenced to confirm sequence integrity. For protein overexpression in *Yersinia*, desired genes were cloned into the pBAD/myc-HIS A vector under the control of an arabinose-inducible promoter. For protein complementation/overexpression in *Bartonella*, selected genes were cloned into plasmid *pBZ485_empty* under the control of a taclac promoter. For the generation of bacterial knockouts, the previously described two-step gene replacement procedure was used (2). The sequences of all oligonucleotide primers used in this study are listed in supplementary Table S2. A detailed description for the construction of each plasmid is presented in supplementary Table S3.

Plasmids were introduced into *Bartonella* strains by conjugation from *E. coli* strain MFDpir using two-parental mating (3). Plasmids were introduced into electro-competent *Yersinia* by electroporation as previously described (4).

##### Infection of RAW macrophages with *Bartonella*

Bartonellae were cultured as described above. One day before infection, RAW 264.7 macrophages were seeded at the following densities: 5 x $10^5$ cells per well in a 12-well plate or 1 x $10^6$ cells in a 6-well plate. RAW LgBiT macrophages were seeded in white 96-well plates (Corning, Cat. CLS3610) at a density of 1 x $10^4$ cells per well. If required, bacterial cultures were supplemented with 100 µM IPTG 1 h prior to infection to induce protein expression. Cell culture medium was replaced with infection medium (DMEM Glutamax, supplemented with 1% FCS). Cells were infected at a multiplicity of infection (MOI) of 50, if not stated otherwise. To synchronize bacterial attachment, plates were centrifuged at 500 g for 3 min. Infected cells were incubated at 5% $CO_2$ and 37°C for indicated time periods. If indicated, cells were stimulated with 100 ng/ml LPS (lipopolysaccharides from *E. coli* O26:B6, Sigma-Aldrich) during the last hour of infection. Supernatants were analyzed by Ready-SET-Go! ELISA kits for TNF-α secretion (Thermo Fisher Scientific). Adherent cells were harvested, lysed and analyzed by immunoblot. For monitoring effector injection, cells were washed with PBS and supplemented with the Nano-Glo® Live Cell Reagent (Promega, Cat. N2011). Luminescence reading was carried out in the Synergy H4 plate reader (BioTek).

**Infection of RAW LgBiT macrophages with *Yersinia***

*Yersinia* strains were grown as described previously. 1 x $10^4$ RAW LgBiT macrophages per well were seeded in white 96-well plates. 2 h prior to infection, *Yersinia* were shifted into a 37°C water bath shaker and supplemented with 0.2% arabinose to induce protein expression. Bacteria were washed twice with infection medium (DMEM supplemented with 1% FCS). Cells were infected with indicated MOIs and bacterial attachment synchronized by centrifugation (500 g, 3 min). After 2 h, cells were washed with PBS and supplemented with the Nano-Glo® Live Cell Reagent (Promega, Cat. N2011). Luminescent signal was measured using the Synergy H4 plate reader.

**Transient transfection of HeLa cells and determination of ByeA targets**

To generate HeLa cells transiently expressing the ByeA wild-type or $ByeA^{C194A}$ mutant of *B. taylorii*, transient transfection was performed as described previously (5, 6). In brief, 3 x $10^5$ HeLa cells per well were seeded in a 6-well plate and transfected with a total of 1 µg plasmid DNA following the FuGENE transfection protocol (FuGENE® HD Transfection Reagent, Promega, Cat. E2311). After 24 h of transfection, cell culture medium was replaced with DMEM supplemented with 20 mM HEPES for 12-16 h for serum starvation. To trigger the activation of MAPK and NF-κB signaling pathways, HeLa cells were stimulated with 20 ng/ml human recombinant TNF-α (BioLegend, Cat. 570102). After the indicated incubation time, immunoblot analysis was performed.

**NanoLuc-based effector translocation assay**

To assess the capacity of the HiBiT-FLAG fragment or effectors fused to HiBiT-FLAG to complement LgBiT to a functional luciferase, we used the Nano-Glo® HiBiT Lytic Detection System (Promega, Cat. N3030). To determine the linear range of the luminescence signal we performed a dilution series with different numbers of bacteria. 1 x $10^7$, 1 x $10^8$, 5 x $10^8$ or 1 x $10^9$

bacteria were washed in PBS and resuspended in 100 µL PBS. 100 µL LSC buffer containing the substrate (1:50 v/v) and the LgBiT protein (1:100 v/v) were added and the luminescent signal measured using the Synergy H4 plate reader. Accordingly, 1 x $10^8$ bacteria were used to test functionality of the HiBiT-fused effectors if not indicated otherwise.

Effector translocation into RAW LgBiT macrophages was quantified by measuring the luminescent signal using the Synergy H4 plate reader. RAW LgBiT macrophages were infected with *Bartonella* or *Yersinia* as described previously. Cell culture supernatant was aspirated and cells were gently washed in pre-warmed PBS. The Nano-Glo® Live Cell Reagent (Promega, Cat. N2011) was prepared as instructed by manufacturer. A final volume of 100 µL PBS per well was supplemented with 25 µL of the Nano-Glo® Live Cell assay buffer containing the substrate. The following plate reader settings were used: temperature 37°C, shaking sequence 30 sec at 300-500 rpm, delay 10 min, autoscale, integration time 5 sec.

**SDS-Page and Immunoblot analysis**

SDS-PAGE and immunoblotting were performed as described (7). To verify protein expression levels, cells were collected, washed in ice-cold PBS and lysed by adding Novagen's PhosphoSafe Extraction Buffer (Merck, Cat. 71296-3) complemented with cOmplete™ Mini EDTA-free Protease Inhibitor Cocktail (Roche, Cat. 11836170001). Protein concentrations were quantified using the Pierce BCA Protein Assay Kit (Thermo Fisher Scientific, Cat. 23225). Lysates were mixed with 5X SDS sample buffer and resolved on 4–20% Mini-PROTEAN® TGX™ Precast Protein Gels (BioRad, Cat. 4561093). Pre-stained Precision Plus Protein Dual Color Standard (BioRad, Cat. 1610374) was used as protein size reference. Proteins were transferred onto Amersham Hybond® PVDF membrane (0.2 µm or 0.45 µm pore size). Membranes were probed with primary antibodies directed against the protein of interest: α-FLAG (Sigma-Aldrich, Cat. F1804), α-c-myc (Roche, Cat. 11667149001), α-p38 (Cell Signaling Technology, Cat. 8690), α-p-p38 (T180/Y182) (Cell Signaling Technology, Cat. 4511), α-JNK (Cell Signaling Technology, Cat. 9252), α-p-JNK (T183/Y185) (Cell Signaling Technology, Cat. 4668.), α-ERK1/2 (Cell Signaling Technology, Cat. 9102), α-p-ERK1/2 (T202/Y204) (Cell Signaling Technology, Cat. 9101), α-p65 (Cell Signaling Technology, Cat. 8242), α-p-p65 (S536) (Cell Signaling Technology, Cat. 3033). The detection of ByeA was performed using a polyclonal antibody targeting YopP of *Y. enterocolitica* (provided by T3 Pharmaceuticals, No. 95). Detection was performed with horseradish peroxidase-conjugated antibodies directed against rabbit or mouse IgG (HRP-linked α-mouse IgG (Cell Signaling Technology, Cat. 7076), HRP-linked α-rabbit IgG (Cell Signaling Technology, Cat. 7074)). Immunoblots were developed using LumiGLO® Peroxidase Chemiluminescent Substrate Kit (Seracare, Cat. 5430-0040) and imaged using the Fusion FX device (Vilber). If required, images were adjusted in brightness and contrast using the ImageJ software.

#### Quantification of secreted TNF-α

TNF-α was quantified in cell culture supernatants of infected RAW macrophages by Ready-SET-Go! ELISA Kit (ThermoFisher, Cat. 88-7324-88). 96-well assay plates (Costar, Cat. 9018) were coated overnight at 4°C with capture antibody in coating buffer. Afterwards the plates were washed with wash buffer (PBS containing 0.05% Tween-20). Unspecific binding was blocked by adding assay diluent (provided in the kit) to each well at RT for 1 h. Before adding the samples, one wash step was performed. The samples were pre-diluted 1:6 in assay diluent. Subsequently, samples were diluted twice by serial 2-fold dilutions on the plate. The respective lyophilized standard was resolved as requested, added to the plate and diluted by serial 2-fold dilutions. The plate was incubated at 4°C overnight. 5 wash steps were performed. The respective detection antibody was added and the plate incubated for 1 hour at RT. After another 5 wash steps, horseradish peroxidase-conjugated avidin was added for 30 min at RT. After 7 washes ELISA substrate solution was added for 5 to 15 min at RT and the reaction stopped by adding 1M $H_3PO_4$. Absorbance was read at 450 and 570 nm.

#### Identification of YopJ homologs in the *Bartonella* genomes

Candidates of YopJ homologs in *Bartonella* were identified by BLASTP searches using YopP of *Y. enterocolitica* as query sequence. Identical duplicates were excluded. A set of 34 candidates were recovered with up to four different YopJ copies in the same *Bartonella* strain. No potential homologs could be identified in the lineage 4 strains *B. henselae*, *B. koehlerae* and *B. birtlesii*. YopJ sequences of the following strains were used in this analysis with protein accession numbers shown in brackets: *B. sp. 1-1C* (UniProt: E6YU28, E6YU30, E6YW76, E6YW79), *B. alsatica* IBS382 (UniProt: J1IT78), B*. clarridgeiae CIP 104772 / 73* (UniProt: E6YIW6, E6YGE3, E6YIF4, E6YIF0), *B. doshiae NCTC 12862* (UniProt: A0A380ZBF3), *B. elizabethea F9251* (UniProt: J0RD58), *B. grahamii as4aup* (UniProt: C6AAE6, C6AAE7), *B. quintana Toulousse* (UniProt: A0A0H3LWL4), *B. rattimassilensis 15908* (UniProt: J1JSC6), *B. rochalimae ATCC BAA-1498* (UniProt: E6YMH5, E6YMI1, E6YKC5, E6YKC3), *B. sp. AR 15-3* (UniProt: E6YRW6, E6YPL1, E6YPL0), *B. sp. JB15* (UniProt: A0A1S6XI99, A0A1S6XIC2, A0A1S6XG80, A0A1S6XI49), *B. taylorii* IBS296 (same sequence as *B. taylorii* 8TBB, UniProt: J1KJI9), *B. tribocorum CIP 105476 / IBS506* (UniProt: A9IXZ4, AI9XZ6, AI9XZ8), *B. vinsonii subsp. arupensis OK-94-513* (UniProt: J1JVH0), *B. vinsonii subsp. berkhoffi* Tweed (UniProt: N6UZ35), *B. washoeensis 085-0475* (UniProt: J1JGP6, J0QL81).

#### Phylogenetic analysis of YopJ homologs and homology modeling of ByeA and YopP

Protein sequences were aligned using ClustalW implemented in Geneious Prime 2019.0.4 using standard settings. The alignment was manually curated to remove largely gapped sequences. Maximum likelihood phylogenies were constructed using PhyML implemented in Geneious with standard settings and bootstrap 100. YopP of *Y. enterocolitica* and AvrA of *Salmonella*

Typhimurium were chosen as outgroups. The structure of the AvrA$^{\Delta1L140}$-IP6-CoA complex (PDB: 6BE0) served as input structure for the homology modeling. The superimposition was carried out using the align-algorithm implemented in PyMOL (version 2.3.4). The electrostatic potential was created using the "APBS Tools2.1" plugin.

**Synteny analysis in L3 and L4 *Bartonella* species**

Synteny analysis of *byeA* was inferred with SyntTax (8) with default search parameters. Obtained results were confirmed by genome alignment using the MAUVE plugin in Geneious Prime v2019.0.4. The database given in SyntTax was supported by a database generated in Geneious from annotated *Bartonella* species (SI Appendix, Table S4).

#### Supplementary figures

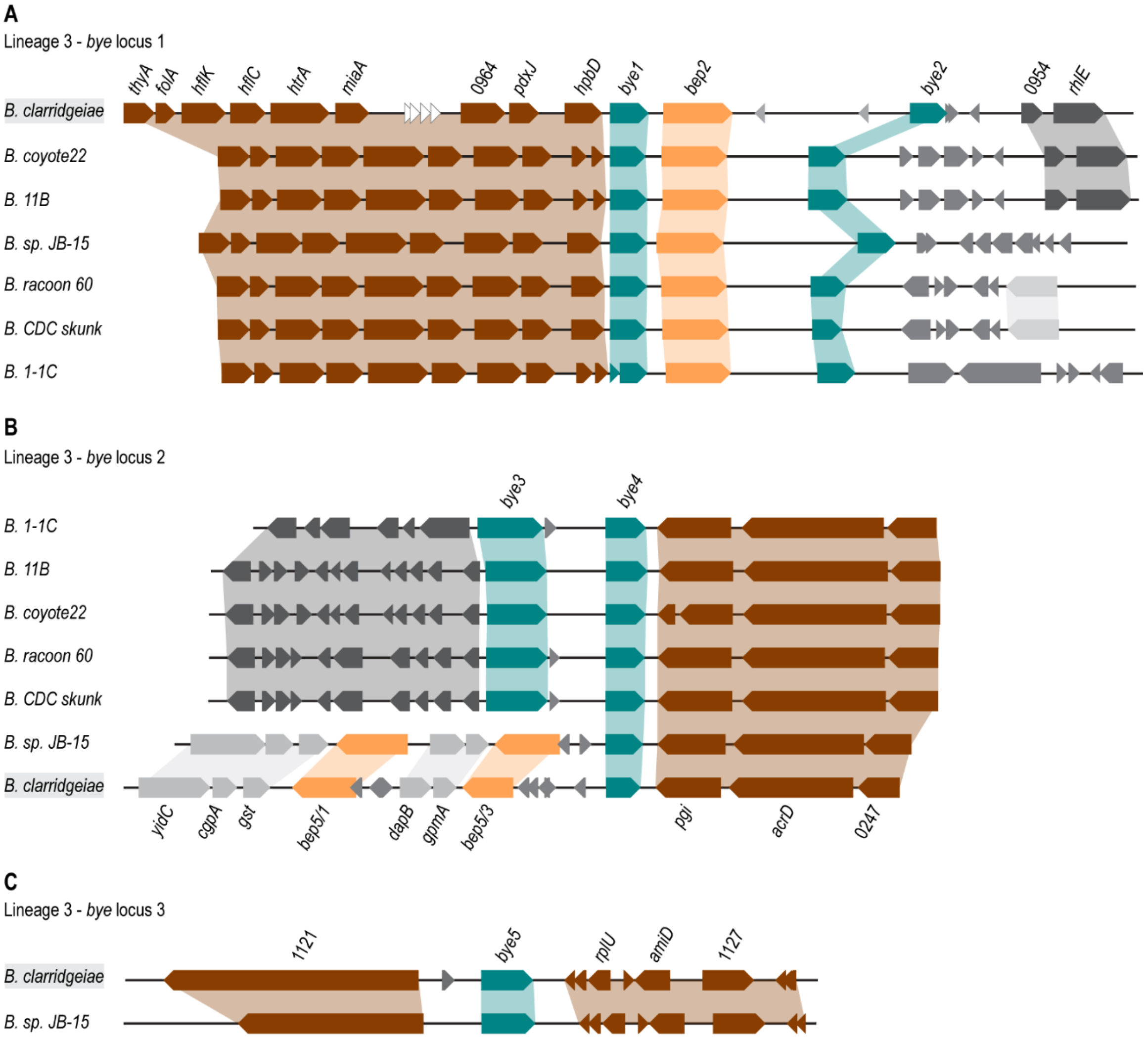


**Fig. S1. Genomic regions encoding different *bye* homologs present in *Bartonella* species of lineage 3.** The genomic regions containing the *bye* loci are compared between seven *Bartonella* species of lineage 3 (*B. clarridgeiae*, *B. coyote22*, *B. 11B*, *B. sp. JB-15*, *B. raccoon 60*, *B. CDC skunk* and *B. 1-1C*). Depending on the species, up to three loci encoding *bye* homologs were identified. Genes are depicted as arrows using the following color-code: *bye* homologs (teal), *bep* homologs (orange), identified gene homologs present in all compared species (brown) and gene homologs only partially present in different species (grey). Gene identifiers shown for *B. clarridgeiae* (highlighted in grey). **(A)** locus containing *bye1* and *bye2*, which are in close proximity to *bep2*. **(B)** locus containing *bye 3* and *bye4*. *B. clarridgeiae* and *B. sp. JB-15*, which encode *bye4* in close proximity to *bep5* homologs seem to lack *bye3*. These two species encode *bye5* in the third locus identified, shown in **(C)**.

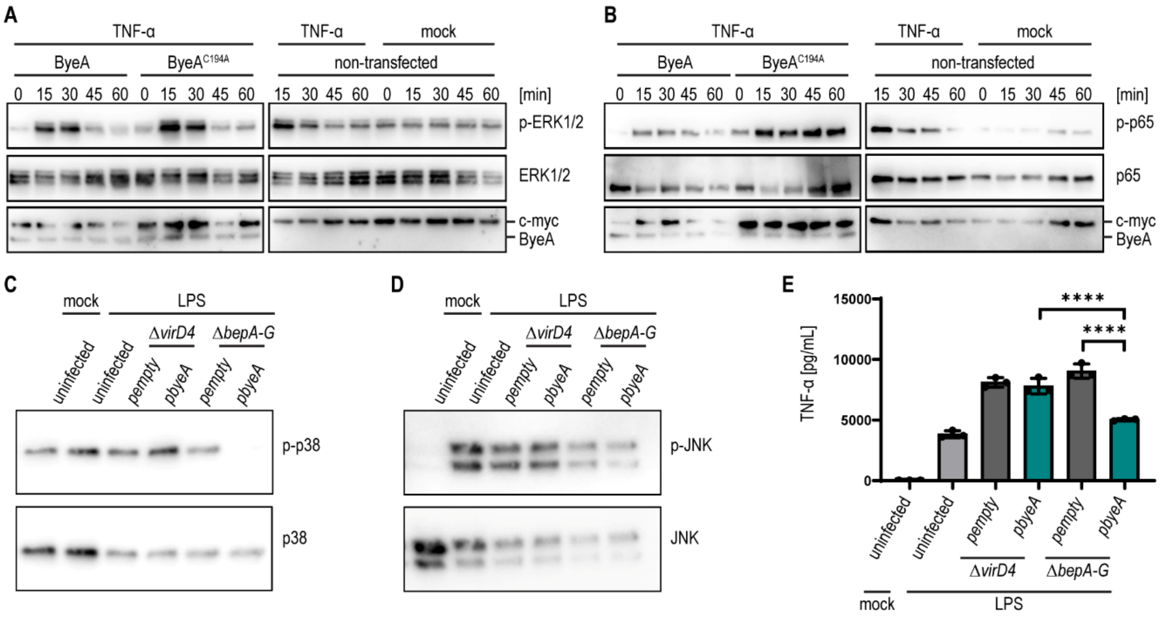


**Fig. S2. ByeA inhibits phosphorylation of p38 and JNK. (A, B**) HeLa cells were transiently transfected with expression constructs for ByeA wild-type or the catalytically-inactive ByeA$^{C194A}$ mutant carrying an N-terminal c-myc epitope tag. Following 12-16 h of serum starvation, cells were treated with 20 ng/ml human recombinant TNF-α for the indicated times. HeLa cells were harvested, lysed and analyzed by immunoblot using the indicated antibodies. Expression levels of ByeA constructs were monitored with an anti-c-myc tag antibody. **(A)** Phosphorylation of ERK or total ERK levels were monitored using the monoclonal antibodies against p-ERK1/2 (T202/Y204) or ERK, respectively. **(B)** Phosphorylation of p65 or total p65 levels were monitored using the monoclonal antibodies against p-p65 (S536) and p65, respectively. **(C-E)** RAW 264.7 macrophages were infected for 24 h at MOI 50 with *B. henselae* wild-type, the Δ*virD4* or Δ*bepA-G* mutants expressing ByeA from a plasmid (p*byeA*) or containing the empty plasmid (p*empty*). Except for the untreated control, cells were co-stimulated with 100 ng/ml *E. coli* LPS during the last hour of infection. **(C)** Immunoblot analysis using monoclonal antibodies against p-p38 (T180/Y182) and p38. **(D)** Immunoblot analysis using monoclonal antibodies against p-JNK (T183/Y185) and JNK. **(E)** Secreted TNF-α was quantified by ELISA. All data show representative results from three independent experiments. **(E)** shows the mean ±SD of technical triplicates. Data was analyzed using one-way ANOVA with multiple comparisons (Tukey's multiple comparison test), **** $p < 0.0001$.

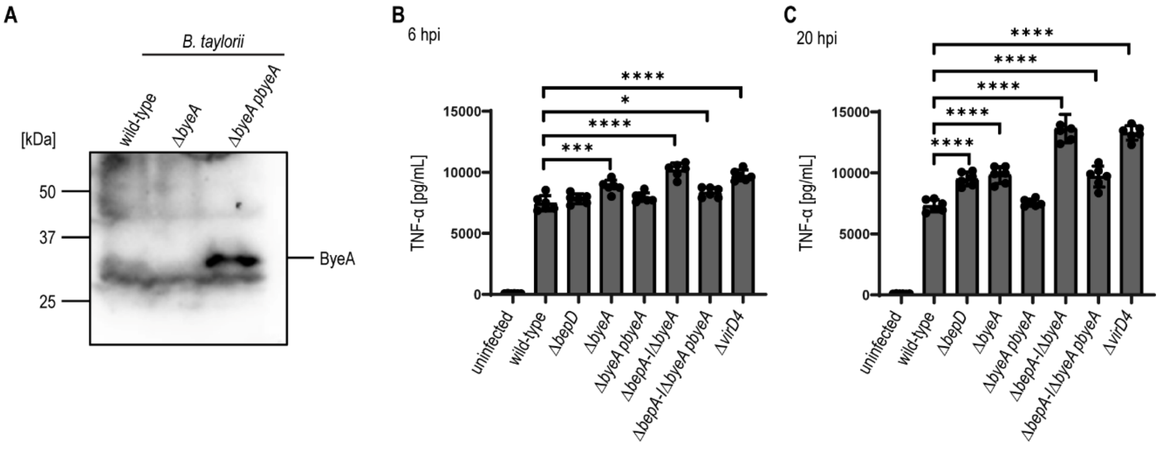


**Fig. S3. ByeA downregulates the pro-inflammatory TNF-α secretion in a VirB/VirD4 T4SS-dependent manner.** *B. taylorii* Δ*virD4* or Δ*bepA-I*Δ*byeA* mutants were incubated for 24 h in M199 + 10% FCS at 28°C and 5% $CO_2$. To induce protein expression, the cultures were supplemented with 100 µM IPTG for at least 60 min prior to infection. **(A)** Expression of endogenous ByeA and ByeA from plasmid in *B. taylorii* was investigated by immunoblot using polyclonal antibody against YopP of *Y. enterocolitica*. *B. taylorii* Δ*byeA* served as control. Estimated molecular mass of ByeA is 34.3 kDa. **(B, C)** RAW 264.7 macrophages were infected at MOI 50 with *B. taylorii* wild-type, the translocation-deficient mutant Δ*virD4*, mutants lacking either *bepD* or *byeA*, mutants depleted of all *beps* and *byeA* (Δ*bepA-I*Δ*byeA*) or deletion mutants expressing ByeA under the control of an IPTG-inducible promoter. **(B)** 6 hpi and **(C)** 20 hpi secreted TNF-α levels were assessed by ELISA. All data show representative results from three independent experiments. **(B, C)** show the mean ±SD of 6 technical replicates. Data was analyzed using one-way ANOVA with multiple comparisons (Tukey's multiple comparison test), * $p < 0.05$, *** $p < 0.001$, **** $p < 0.0001$.

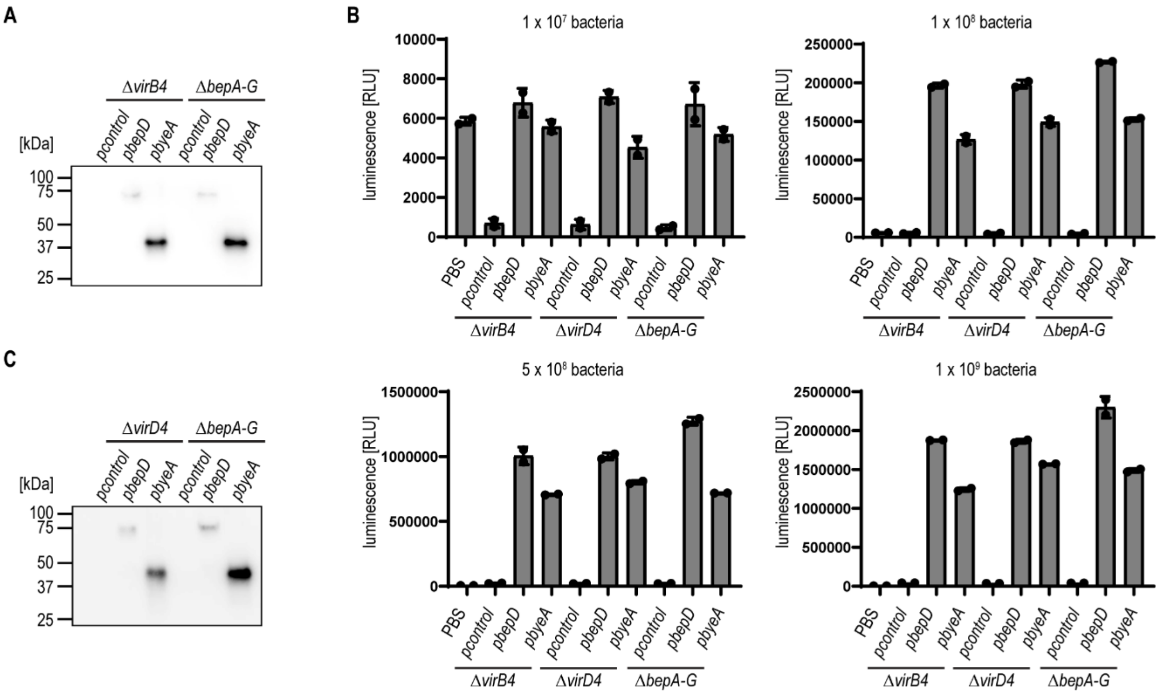


**Fig. S4. ByeA is translocated in a VirB4- and VirD4-dependent manner.** *B. henselae* Δ*virB4,* Δ*virD4* or Δ*bepA-G* mutants were incubated for 24 h in M199 + 10% FCS at 35°C and 5% $CO_2$. To induce protein expression, the cultures were supplemented with 100 µM IPTG for at least 60 min prior to infection. For simplicity, the sequence *hibit-flag*, which is N-terminally fused to all depicted effectors in *Bartonella* and present in the empty vector, is not shown in the plasmid names. **(A)** Expression of HiBiT-FLAG, HiBiT-FLAG-BepD (*bepD* sequence of *B. henselae*) and HiBiT-FLAG-ByeA in Δ*virB4* or Δ*bepA-G* mutants was assessed by immunoblot using an antibody against FLAG-epitope tag. **(B)** The interaction of HiBiT-FLAG, HiBiT-FLAG-BepD and HiBiT-FLAG-ByeA with LgBiT was tested using the Nano-Glo® HiBiT Lytic Detection System. Indicated numbers of lysed bacteria (expression of fusion proteins shown in **(A and C)**) were supplemented with purified LgBiT and substrate. **(C)** Expression of HiBiT-FLAG, HiBiT-FLAG-BepD and HiBiT-FLAG-ByeA in Δ*virD4* or Δ*bepA-G* mutants was assessed by immunoblot using an antibody against FLAG-epitope tag. All data show representative results from three independent experiments. **(B)** shows the mean ±SD of technical duplicates.

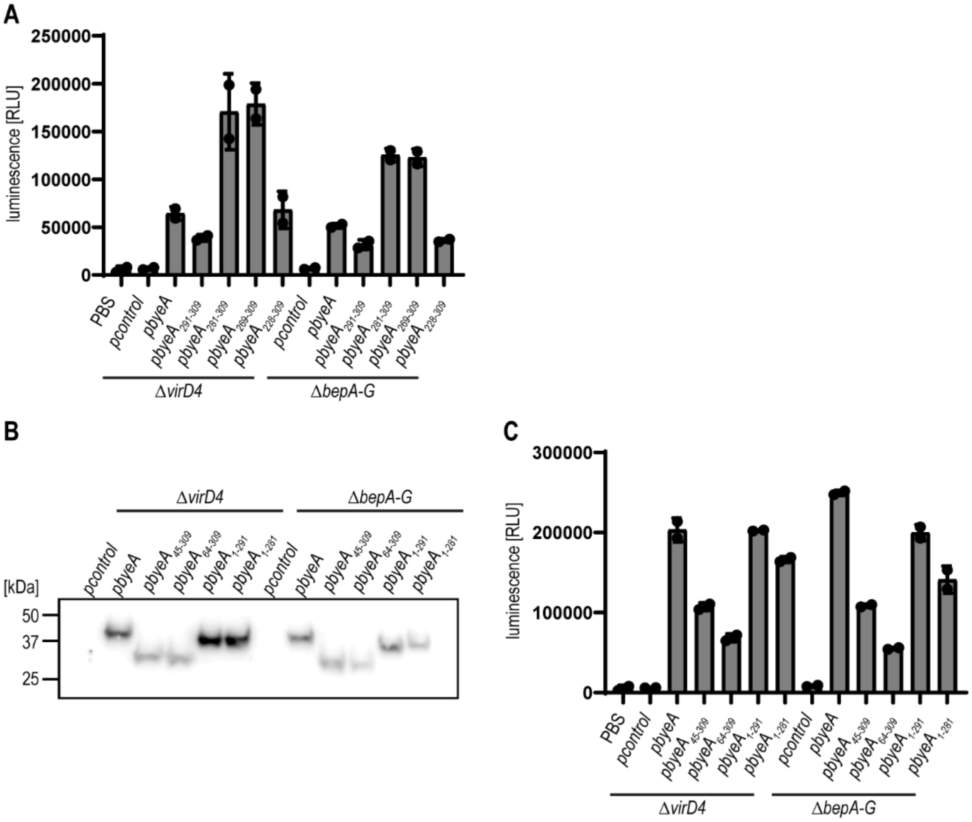


**Fig. S5. Translocation of ByeA via the VirB/VirD4 T4SS depends on N-proximal helix and the C-terminus.** *B. henselae* Δ*virD4* or Δ*bepA-G* mutants were incubated for 24 h in M199 + 10% FCS at 35°C and 5% $CO_2$. To induce protein expression, the cultures were supplemented with 100 µM IPTG for at least 60 min prior to infection. For simplicity, the sequence *hibit-flag*, which is N-terminally fused to all depicted effectors in *Bartonella*, is not shown in the plasmid names. The expression of different effectors fused to HiBiT-FLAG was assessed by immunoblot using an antibody against FLAG-epitope tag. The interaction of effectors fused to HiBiT-FLAG with LgBiT was tested using the Nano-Glo® HiBiT Lytic Detection System. Lysed bacteria were supplemented with purified LgBiT and substrate. **(A)** The interaction of HiBiT-FLAG-ByeA, HiBiT-FLAG-ByeA$_{291-309}$, HiBiT-FLAG-ByeA$_{281-309}$, HiBiT-FLAG-ByeA$_{269-309}$ and HiBiT-FLAG-ByeA$_{228-309}$ with LgBiT in Δ*virD4* or Δ*bepA-G* mutants. **(B)** Expression of HiBiT-FLAG-ByeA, HiBiT-FLAG-ByeA$_{45-309}$, HiBiT-FLAG-ByeA$_{64-309}$, HiBiT-FLAG-ByeA$_{1-291}$ and HiBiT-FLAG-ByeA$_{1-281}$ in Δ*virD4* or Δ*bepA-G* mutants. **(C)** Interaction of effectors shown in **(B)** with LgBiT. All data show representative results from three independent experiments. **(A, C)** show the mean ±SD of technical duplicates.

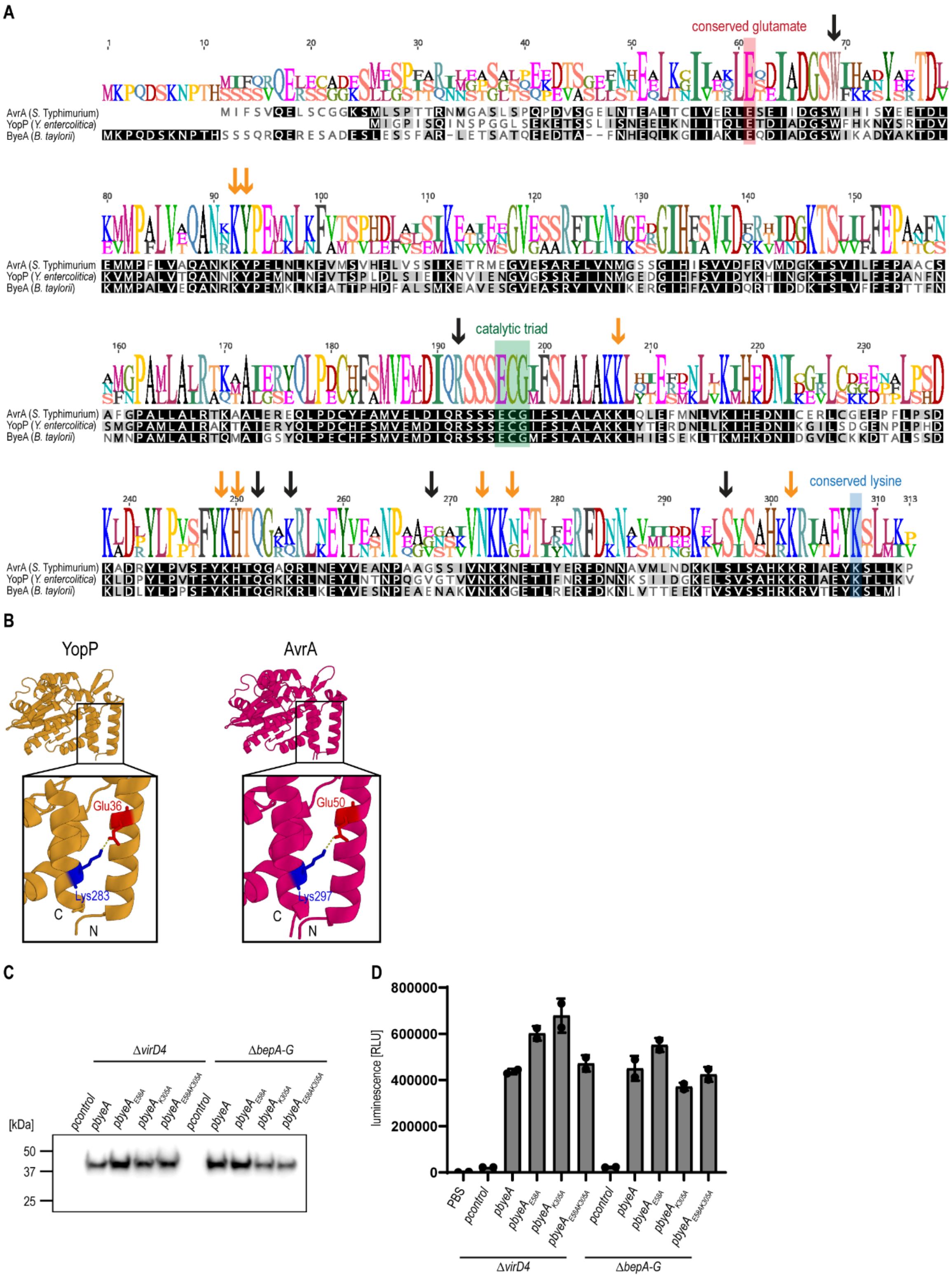


**Fig. S6. Conserved charged amino acids in N- and C-terminus are important for the translocation via the T4SS in *Bartonella.* (A)** Sequence alignment of ByeA of *B. taylorii*, YopP of *Y. enterocolitica* and AvrA of *S.* Typhimurium. Amino acids were highlighted in grey scale if all amino acids at a given position are 100 % conserved (black), 80-100% similar (dark grey), or 60-

80 % similar (light grey) based on Blosum62 score matrix with a threshold of 1. The catalytic triad of His, Glu and Cys residues is highlighted in green. The conserved glutamate residue (highlighted in red) and lysine residue (highlighted in blue) are important for the translocation via the VirB/VirD4 T4SS. The binding sites for IP6 are indicated as orange arrows, the binding sites for acetyl-CoA as black arrows. **(B)** Structures of YopJ homologs were modeled using PyMOL with focus on the C-terminal and N-terminal helices. Positively charged amino acids are depicted in blue, negatively charged amino acids shown in red. Predicted salt bridges are shown in yellow. Predicted interactions are indicated (YopP: E36 and K283, AvrA: E50 and K297). **(C)** *B. henselae* Δ*virD4* or Δ*bepA-G* mutants were incubated for 24 h in M199 + 10% FCS at 35°C and 5% $CO_2$. To induce protein expression, the cultures were supplemented with 100 µM IPTG for at least 60 min prior to infection. For simplicity, the sequence *hibit-flag*, which is N-terminally fused to all depicted effectors in *Bartonella*, is not shown in the plasmid names. Expression of HiBiT-FLAG-ByeA, HiBiT-FLAG-ByeA$_{E58A}$, HiBiT-FLAG-ByeA$_{K305A}$ or HiBiT-FLAG-ByeA$_{E58AK305A}$ in Δ*virD4* or Δ*bepA-G* mutants was assessed by immunoblot using an antibody against FLAG-epitope tag. **(D)** The interaction of effectors shown in **(C)** with LgBiT was tested using the Nano-Glo® HiBiT Lytic Detection System. Lysed bacteria were supplemented with the purified LgBiT protein and the substrate. All data show representative results from three independent experiments. **(D)** shows the mean ±SD of technical duplicates.

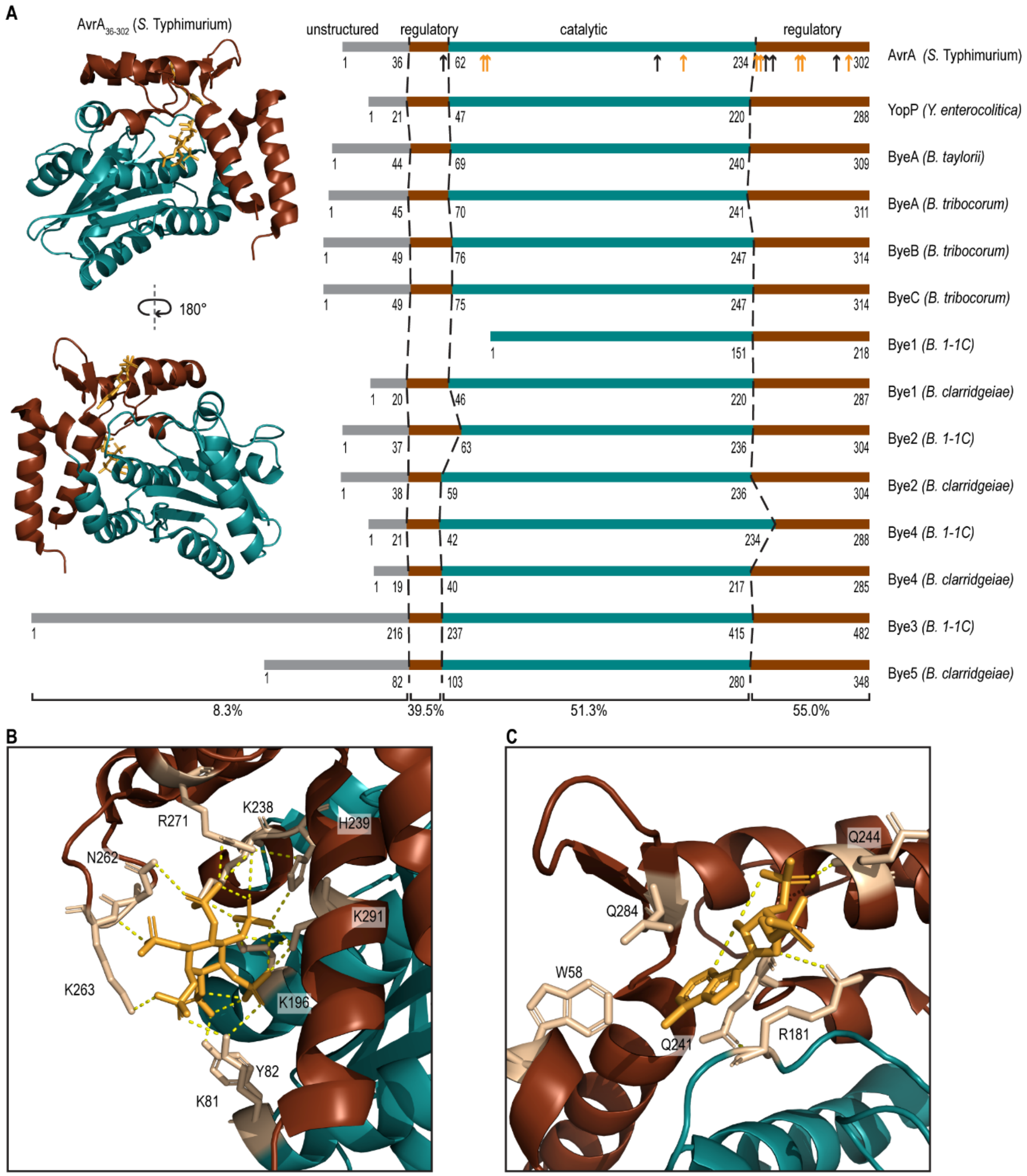


**Fig. S7. Regulatory and catalytic domains present in Bye homologs of L3 and L4 bacteria.** **(A)** The unstructured N-terminus (grey), the regulatory domains (brown) and catalytic core (teal) of AvrA were identified previously (9). The structure of AvrA is shown in complex with IP6 (top) and acetyl-CoA (bottom). The binding sites for IP6 are indicated as orange arrows, the binding sites for acetyl-CoA as black arrows. According to the domain boundaries of AvrA, we identified the regulatory and catalytic domains for Bye homologs present in L3 and L4 Bartonellae. **(B, C)** Residues present in the regulatory and catalytic domain of AvrA important for **(B)** the IP6 interaction or **(C)** the acetyl-CoA interaction.

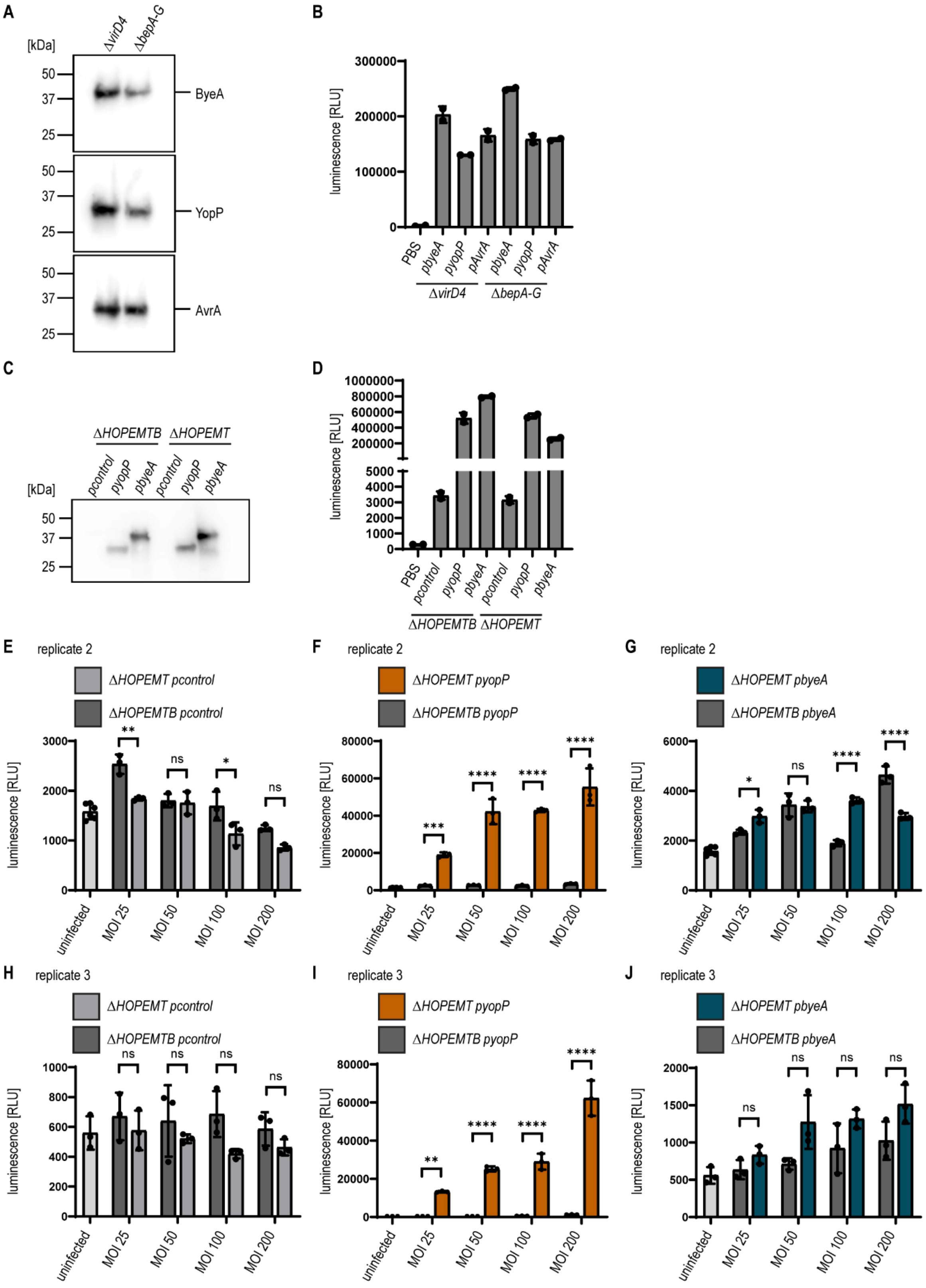


**Fig. S8. Translocation of YopJ homologs through T3SS and VirB/VirD4 T4SS. (A, B)** *B. henselae* Δ*virD4* or Δ*bepA-G* mutants were incubated for 24 h in M199 + 10% FCS at 35°C and

5% $CO_2$. To induce protein expression, the cultures were supplemented with 100 µM IPTG for at least 60 min prior to infection. For simplicity, the sequence *hibit-flag*, which is N-terminally fused to all depicted effectors in *Bartonella*, is not shown in the plasmid names. **(A)** Expression of HiBiT-FLAG-ByeA, HiBiT-FLAG-YopP or HiBiT-FLAG-AvrA in Δ*virD4* or Δ*bepA-G* mutants was assessed by immunoblot using an antibody against FLAG-epitope tag. **(B)** The interaction of HiBiT-FLAG-ByeA, HiBiT-FLAG-YopP or HiBiT-FLAG-AvrA with LgBiT was tested using the Nano-Glo® HiBiT Lytic Detection System. Lysed bacteria (expression of fusion proteins shown in **(A)**) were supplemented with LgBiT protein and substrate. **(C-J)** *Y. enterocolitica* effector-less mutant Δ*HOPEMT* and the translocation-deficient mutant Δ*HOPEMTB* were supplemented with 0.2% arabinose to induce protein expression. For simplicity, the *flag-hibit* sequence, which is C-terminally fused to all depicted effectors in *Yersinia*, is not indicated in the plasmid names. **(C)** Expression of YopP-FLAG-HiBiT and ByeA-FLAG-HiBiT was validated by immunoblot using specific antibody against the FLAG-epitope tag. **(D)** The interaction of FLAG-HiBiT, YopP-FLAG-HiBiT and ByeA-FLAG-HiBiT with LgBiT was tested using the Nano-Glo® HiBiT Lytic Detection System. Lysed bacteria (expression of fusion proteins shown in **(C)**) were supplemented with the purified LgBiT protein and the substrate. **(E-J)** RAW LgBiT macrophages were infected using the depicted MOIs for 2 h with **(E, H)** *Y. enterocolitica ΔHOPEMT* or *ΔHOPEMTB* expressing FLAG-HiBiT (p*control*), **(F, I)** *Y. enterocolitica ΔHOPEMT* or *ΔHOPEMTB* expressing YopP-FLAG-HiBiT (p*yopP*) or **(G, J)** *Y. enterocolitica ΔHOPEMT* or *ΔHOPEMTB* expressing ByeA-FLAG-HiBiT (p*byeA*). **(B, D)** show the mean ±SD of technical duplicates and **(E-J)** of technical triplicates. The luminescence of the complemented split NanoLuc was measured. Data was analyzed using one-way ANOVA with multiple comparisons (Tukey's multiple comparison test), ns = not significant, * $p < 0.05$, ** $p < 0.01$, *** $p < 0.001$ **** $p < 0.0001$. All data show representative results from three independent experiments.

**Supplementary Tables**

**Table S1.** List and construction of all bacterial strains of this study

| Strain | Genotype | Reference/Source | Identifier / Description / Construction |
|---|---|---|---|
| ***Escherichia coli*** | | | |
| Novablue | *endA1 hsdR17* ($r_{K12}^{-}$ $m_{K12}^{+}$) *supE44 thi-1 recA1 gyrA96 relA1 lac F'[proA⁺B⁺ lacI*$^{q}$*ZΔM15::Tn10]* | Novagen | Standard cloning strain |
| HST08 | *F-, endA1, supE44, thi-1, recA1, relA1, gyrA96, phoA, Φ80d lacZΔ M15, Δ (lacZYA - argF) U169, Δ (mrr - hsdRMS - mcrBC), ΔmcrA, λ-* | Takara | Standard cloning strain |
| JKE201 | MFDpir Δ*mcrA* Δ*(mrr-hsdRMS-mcrBC) aac(3)IV::lacI*$^{q}$ | (3) | derivative of MFDpir lacking EcoKI, the three type IV restriction systems, restored gentamicin sensitivity, harboring *lacI*$^{q}$ allele |
| ***Bartonella henselae*** | | | |
| *B. henselae* Houston-1 | *rpsL* | (10) | RSE247, spontaneous SmR strain of *B. henselae* ATCC49882T, serving as wild-type |
| | *rpsL* Δ*virD4* | (7) | GS0221; virD4 deletion mutant, derivative of RSE247 |
| | *rpsL* Δ*virD4 / pHiBiT-FLAG* | (11) | KFB286; GS0221 containing pKF059 |
| | *rpsL* Δ*virD4 / pHiBiT-FLAG-bepD*$_{Bhe}$ | (11) | KFB211; GS0221 containing pKF027 |
| | *rpsL* Δ*virD4 / pHiBiT-FLAG-yopP* | This study | KFB219; GS0221 containing pKF029 |
| | *rpsL* Δ*virD4 / pHiBiT-FLAG-avrA* | This study | KFB317; GS0221 containing pKF063 |
| | *rpsL* Δ*virD4 / pHiBiT-FLAG-byeA* | This study | KFB213; GS0221 containing pKF028 |
| | *rpsL* Δ*virD4 / pHiBiT-FLAG-yopJ*$_{45\text{-}309}$ | This study | KFB253; GS0221 containing pKF042 |
| | *rpsL* Δ*virD4 / pHiBiT-FLAG-byeA*$_{64\text{-}309}$ | This study | KFB315; GS0221 containing pKF062 |
| | *rpsL* Δ*virD4 / pHiBiT-FLAG-byeA*$_{1\text{-}291}$ | This study | KFB255; GS0221 containing pKF043 |
| | *rpsL* Δ*virD4 / pHiBiT-FLAG-byeA*$_{1\text{-}281}$ | This study | KFB257; GS0221 containing pKF044 |
| | *rpsL* Δ*virD4 / pHiBiT-FLAG-byeA*$_{292\text{-}309}$ | This study | KFB281; GS0221 containing pKF055 |
| | *rpsL* Δ*virD4 / pHiBiT-FLAG-byeA*$_{282\text{-}309}$ | This study | KFB283; GS0221 containing pKF056 |
| | *rpsL* Δ*virD4 / pHiBiT-FLAG-byeA*$_{269\text{-}309}$ | This study | KFB297; GS0221 containing pKF057 |
| | *rpsL* Δ*virD4 / pHiBiT-FLAG-byeA*$_{228\text{-}309}$ | This study | KFB299; GS0221 containing pKF058 |

| | | | |
|---|---|---|---|
| | *rpsL ΔvirD4 / pHiBiT-FLAG-byeA$_{K305A}$* | This study | KFB330; GS0221 containing pKF051 |
| | *rpsL ΔvirD4 / pHiBiT-FLAG-byeA$_{E58A}$* | This study | KFB332; GS0221 containing pKF065 |
| | *rpsL ΔvirD4 / pHiBiT-FLAG-byeA$_{E58AK305A}$* | This study | KFB334; GS0221 containing pKF066 |
| | *rpsL ΔbepA-G* | (7) | MSE150; *bepA-bepG* deletion mutant, derivative of RSE247 |
| | *rpsL ΔbepA-G / pHiBiT-FLAG* | (11) | KFB276; MSE150 containing pKF059 |
| | *rpsL ΔbepA-G / pHiBiT-FLAG-bepD$_{Bhe}$* | (11) | KFB205; MSE150 containing pKF027 |
| | *rpsL ΔbepA-G / pHiBiT-FLAG-yopP* | This study | KFB217; MSE150 containing pKF029 |
| | *rpsL ΔbepA-G / pHiBiT-FLAG-avrA$_{I}$* | This study | KFB313; MSE150 containing pKF063 |
| | *rpsL ΔbepA-G / pHiBiT-FLAG-byeA* | This study | KFB207; MSE150 containing pKF028 |
| | *rpsL ΔbepA-G / pHiBiT-FLAG-byeA$_{45-309}$* | This study | KFB247; MSE150 containing pKF042 |
| | *rpsL ΔbepA-G / pHiBiT-FLAG-byeA$_{64-309}$* | This study | KFB311; MSE150 containing pKF062 |
| | *rpsL ΔbepA-G / pHiBiT-FLAG-byeA$_{1-291}$* | This study | KFB249; MSE150 containing pKF043 |
| | *rpsL ΔbepA-G / pHiBiT-FLAG-byeA$_{1-281}$* | This study | KFB251; MSE150 containing pKF044 |
| | *rpsL ΔbepA-G / pHiBiT-FLAG-byeA$_{292-309}$* | This study | KFB271; MSE150 containing pKF055 |
| | *rpsL ΔbepA-G / pHiBiT-FLAG-byeA$_{282-309}$* | This study | KFB273; MSE150 containing pKF056 |
| | *rpsL ΔbepA-G / pHiBiT-FLAG-byeA$_{269-309}$* | This study | KFB293; MSE150 containing pKF057 |
| | *rpsL ΔbepA-G / pHiBiT-FLAG-byeA$_{228-309}$* | This study | KFB295; MSE150 containing pKF058 |
| | *rpsL ΔbepA-G / pHiBiT-FLAG-byeA$_{K305A}$* | This study | KFB324; MSE150 containing pKF051 |
| | *rpsL ΔbepA-G / pHiBiT-FLAG-byeA$_{E58A}$* | This study | KFB326; MSE150 containing pKF065 |
| | *rpsL ΔbepA-G / pHiBiT-FLAG-byeA$_{E58AK305A}$* | This study | KFB328; MSE150 containing pKF066 |
| ***Bartonella taylorii*** | | | |
| *B. taylorii IBS296 Sm$^R$* | *rpsL* | (12) | KFB030, spontaneous SmR strain of *B. taylorii IBS296*, serving as wild-type, derivative of LUB046 |
| | *rpsL ΔvirD4* | (11) | KFB146; *virD4* deletion mutant of KFB030 |
| | *rpsL ΔvirD4 / pHiBiT-FLAG* | (11) | KFB291, KFB146 containing pKF059 |
| | *rpsL ΔvirD4 / pHiBiT-FLAG-bepD$_{Bta}$* | (11) | KFB233, KFB146 containing pKF027 |
| | *rpsL ΔvirD4 / pHiBiT-FLAG-byeA* | This study | KFB235; KFB146 containing pKF028 |
| | *rpsL ΔbepD* | (11) | KFB070; *bepD* deletion mutant of KFB030 |

| | | | |
|---|---|---|---|
| | *rpsL ΔbepA-I* | (11) | KFB072; *bepA-bepI* deletion mutant of KFB030 |
| | *rpsL ΔbyeA* | This study | KFB150; *yopJ* deletion mutant of KFB030 |
| | *rpsL ΔbyeA / pbyeA* | This study | ANB0157 ; KFB150 containing pKF017 |
| | *rpsL ΔbepA-IΔbyeA* | This study | KFB154; *yopJ* deletion mutant of KFB072 |
| | *rpsL ΔbepA-IΔbyeA / pbyeA* | This study | ANB0159; KFB154 containing pKF017 |
| | *rpsL ΔbepA-IΔbyeA / pHiBiT-FLAG* | This study | KFB289; KFB154 containing pKF059 |
| | *rpsL ΔbepA-IΔbyeA / pHiBiT-FLAG-*$bepD_{Bta}$ | This study | KFB241; KFB154 containing pKF027 |
| | *rpsL ΔbepA-IΔbyeA / pHiBiT-FLAG-byeA* | This study | KFB243; KFB154 containing pKF028 |
| ***Yersinia enterocolitica*** | | | |
| *Y. enterocolitica* serotype O:9 | strain W22703 (pYV227) | (13) | |
| | *ΔHOPEMTB* | (14) | KFE097; strain W22703 (pYV227) depleted of *yopH*, *yopO*, *yopM*, *yopE*, *yopM*, *yopT* and *yopB* |
| | *ΔHOPEMTB / pFLAG-HiBiT* | This study | KFE119; KFE097 containing pKF060 |
| | *ΔHOPEMTB / pyopP-FLAG-HiBiT* | This study | KFE117; KFE097 containing pKF041 |
| | *ΔHOPEMTB / pbyeA-FLAG-HiBiT* | This study | KFE115; KFE097 containing pKF040 |
| | *ΔHOPEMT* | (15) | KFE096; strain W22703 (pYV227) depleted of *yopH*, *yopO*, *yopM*, *yopE*, *yopM* and *yopT* |
| | *ΔHOPEMT / pFLAG-HiBiT* | This study | KFE113; KFE096 containing pKF060 |
| | *ΔHOPEMT / pyopP-FLAG-HiBiT* | This study | KFE111; KFE096 containing pKF041 |
| | *ΔHOPEMT / pbyeA-FLAG-HiBiT* | This study | KFE109; KFE096 containing pKF040 |

**Table S2.** List of plasmids used in this study

| Plasmid | Backbone | Description | Reference/Source |
|---|---|---|---|
| pTR1000 | | *Bartonella* suicide plasmid with *kanR / rpsL* double-selectable cassette for scarless deletions | (2) |
| pKF009 | pTR1000 | pTR1000 with homology sites to delete *byeA* of *Bartonella taylorii* (homology regions amplified separately and then fused by SOEing PCR) | This study |
| pBZ485 | | new *E. coli* / *Bartonella* shuttle vector based on pCD341 with P*lac*(MQ5); RP4 *oriT* | (16) |
| pKF017 | pBZ485 | Derivative of pBZ485, encodes for *Bta* ByeA fusion protein | This study |
| pKF027 | pBZ485 | Derivative of pBZ485, encodes for HiBiT::FLAG *Bta* BepD fusion protein | (11) |
| pKF028 | pBZ485 | Derivative of pBZ485, encodes for HiBiT::FLAG *Bta* ByeA fusion protein | This study |
| pKF029 | pBZ485 | Derivative of pBZ485, encodes for HiBiT::FLAG *Yer* YopP fusion protein | This study |
| pKF042 | pBZ485 | Derivative of pBZ485, encodes for HiBiT::FLAG *Bta* $ByeA_{45-309}$ fusion protein | This study |
| pKF043 | pBZ485 | Derivative of pBZ485, encodes for HiBiT::FLAG *Bta* $ByeA_{1-291}$ fusion protein | This study |
| pKF044 | pBZ485 | Derivative of pBZ485, encodes for HiBiT::FLAG *Bta* $ByeA_{1-281}$ fusion protein | This study |
| pKF051 | pBZ485 | Derivative of pBZ485, encodes for HiBiT::FLAG *Bta* $ByeA_{K305A}$ fusion protein | This study |
| pKF055 | pBZ485 | Derivative of pBZ485, encodes for HiBiT::FLAG *Bta* $ByeA_{292-309}$ fusion protein | This study |
| pKF056 | pBZ485 | Derivative of pBZ485, encodes for HiBiT::FLAG *Bta* $ByeA_{282-309}$ fusion protein | This study |
| pKF057 | pBZ485 | Derivative of pBZ485, encodes for HiBiT::FLAG *Bta* $ByeA_{269-309}$ fusion protein | This study |
| pKF058 | pBZ485 | Derivative of pBZ485, encodes for HiBiT::FLAG *Bta* $ByeA_{228-309}$ fusion protein | This study |
| pKF059 | pBZ485 | Derivative of pBZ485, encodes for HiBiT::FLAG | This study |
| pKF062 | pBZ485 | Derivative of pBZ485, encodes for HiBiT::FLAG *Bta* $ByeA_{64-309}$ fusion protein | This study |
| pKF063 | pBZ485 | Derivative of pBZ485, encodes for HiBiT::FLAG *Sal* AvrA fusion protein | This study |
| pKF065 | pBZ485 | Derivative of pBZ485, encodes for HiBiT::FLAG *Bta* $ByeA_{E58A}$ fusion protein | This study |
| pKF066 | pBZ485 | Derivative of pBZ485, encodes for HiBiT::FLAG *Bta* $ByeA_{E58AK305A}$ fusion protein | This study |
| pMO006 | pBZ485 | Derivative of pBZ485, encodes for HiBiT::FLAG *Bhe* BepD fusion protein | (11) |
| pKF040 | pBAD | Derivative of pBAD, encodes for FLAG::HiBiT fusion protein | This study |
| pKF041 | pBAD | Derivative of pBAD, encodes for *Bta* ByeA FLAG::HiBiT fusion protein | This study |
| pKF060 | pBAD | Derivative of pBAD, encodes for *Yer* YopP FLAG::HiBiT fusion protein | This study |

**Table S3.** List of oligonucleotide primers used in this study

| Primer | Sequence | Purpose |
|---|---|---|
| prSIM104 | GCACTCCCGTTCTGGATAAT | sequencing pBZ485 insert_fw |
| prAH514 | GGTTTTCCCAGTCACGACG | sequencing pBZ485 insert_rv |
| prKF111 | AGATTACGAATTCCCGGAAGAAGGAGATATACAAATGAAACC GCAAGATTCAAAAAAC | expression *byeA*_fw_BamHI |
| prKF050 | GGCATCAAATTAAGCAGAAGGCCATCCTGACGCCCCCGGGG ATCCAATAGACTAAAATAAAAAGCACAAGGTTATGCTTTTTCT | *ΔbyeA*_US_fw_BamHI |
| prKF051 | AACGTTAAAAATTATATCATCGGTTTCATAATATTTATCTCCGT GTTAAAAGTCG | *ΔbyeA*_US_rv |
| prKF052 | AGATAAATATTATGAAACCGATGATATAATTTTTAACGTTTTTG CCAAGATGTGG | *ΔbyeA*_DS_fw |
| prKF053 | GATGAAACCAAGCTGCAGAGCTTAGCTCTGCAGGTCGACTCT AGAAACCGCGCGTTCAATAGTATTTTCA | *ΔbyeA*_DS_rv_XbaI |
| prKF085 | TACAAACTCACATTTCTCTCCC | sequencing *ΔbyeA*_fw |
| prKF086 | TTGAGCTTTTTTAACAGTGTCG | sequencing *ΔbyeA*_rv |
| prKF087 | TAGGTGATGGTGTTTTGAGG | sequencing *ΔbyeA*_intern |
| prKF112 | GAGTCACTGAATATAAATCTCTCATGATATAAGGTACCTCGAG CGGCCG | expression *byeA*_rv_SalI |
| prKF164 | GAGCCGGGATCCAAGAAGGAGATATACAAATGGTGAG | expression *HiBiT-FLAG*_fw_BamHI |
| prKF165 | ATTCTTTTTTTTGTCATCGTCATCCTTG | expression *HiBiT-FLAG*_rv |
| prKF166 | ATGACAAAAAAAAGAATCATCCATCCCC | expression $bepD_{Bta}$_fw for pKF027 |
| prKF167 | GAGCCGGTCGACTTACATCGCAAAAGCCATTC | expression $bepD_{Bta}$_rv_SalI for pKF027 |
| prKF169 | ATGACAAAAAACCGCAAGATTCAAAAAAC | expression *byeA*_fw for pKF028 |
| prKF170 | GAGCCGGTCGACTTATATCATGAGAGATTTATATTC | expression *byeA*_rv_SalI for pKF028 |
| prKF171 | GAGCCGTCTAGAAAGAAGGAGATATACAAATGGTGAG | expression *FLAG-HiBiT*_fw_XbaI for pKF029 |
| prKF172 | GGCCCAATTTTGTCATCGTCATCCTTG | expression *FLAG-HiBiT*_rv for pKF029 |
| prKF173 | ATGACAAAATTGGGCCAATATCACAAATAAAC | expression *yopP*_fw for pKF029 |
| prKF174 | GAGCCGGTCGACTTATACTTTGAGAAGTGTTTTATATTCAGC | expression *yopP*_rv_SalI for pKF029 |
| prKF176 | GAGCCGACATGTTAAAACCGCAAGATTCAAAAAAC | expression *byeA*_fw_PciI for pKF040 |
| prKF178 | GAGCCGACATGTTAATTGGGCCAATATCACAAATAAAC | expression *yopP*_fw_PciI for pKF041 |
| prKF185 | GAGCCGTCTAGAAAGAAGGAGATATACAAATGCAGCAATCGC ATCAG | expression *FLAG-HiBiT*_rv for pKF028 |
| prKF199 | TTTGTAGTCTATCATGAGAGATTTATATTCAGTGACTC | expression *byeA*_rv for pKF040 |
| prKF200 | TCATGATAGACTACAAAGACCATGACGG | expression *FLAG-HiBiT*_fw for pKF041 |
| prKF201 | GAGCCGGAATTCTTAGCTAATCTTCTTGAACAGCC | expression *FLAG-HiBiT*_rv_EcoRI |
| prKF202 | TTGTAGTCTACTTTGAGAAGTGTTTTATATTCAGCTATTC | expression *yopP*_rv for pKF041 |

| | | |
|---|---|---|
| prKF203 | TCTCAAAGTAGACTACAAAGACCATGACGG | expression *FLAG-HiBiT*_fw for pKF040 |
| prKF205 | ATGATTGAATTTGTCATCGTCATCCTTG | expression *FLAG-HiBiT*_rv for pKF042 |
| prKF206 | ATGACAAATTCAATCATGAACAATTAAAAGG | expression *byeA*$_{45\text{-}309}$_fw for pKF042 |
| prKF207 | GAGCCGGTCGACTTAAGTTTTTTCCTCTGTTGTTACTAAG | expression *byeA*$_{1\text{-}291}$_rv_SalI for pKF043 |
| prKF208 | GAGCCGGTCGACTTAATCGAACCTTTCACGAAG | expression *byeA*$_{1\text{-}291}$_rv_SalI for pKF044 |
| prKF241 | GAGCCGGTCGACTTATATCATGAGAGATTTATATTCAGTGACTCTTTTTCTATGTGATGAAACAGAGACTTTGTCATCGTCATCCTTG | expression *byeA*$_{292\text{-}309}$_rv_SalI for pKF055 |
| prKF242 | GAGCCGGTCGACTTATATCATGAGAGATTTATATTCAGTGACTCTTTTTCTATGTGATGAAACAGAGACAGTTTTTTCCTCTGTTGTTACTAAGTTTTTTTTGTCATCGTCATCCTTG | expression *byeA*$_{282\text{-}309}$_rv_SalI for pKF056 |
| prKF243 | TTTCGCCTTTTTTGTTAACTTTGTCATCGTCATCCTTG | expression *FLAG-HiBiT*_rv for pKF057 |
| prKF244 | AAGGATGACGATGACAAAGTTAACAAAAAAGGCGAAACG | expression *byeA*$_{269\text{-}309}$_fw for pKF057 |
| prKF245 | AGAAGATAAAGCAGTATCTTTGTCATCGTCATCCTTG | expression *FLAG-HiBiT*_rv for pKF058 |
| prKF246 | AAGGATGACGATGACAAAGATACTGCTTTATCTTCTGATAAGTTG | expression *byeA*$_{228\text{-}309}$_fw for pKF058 |
| prKF247 | GAGCCGACATGTTAGACTACAAAGACCATGACGG | expression *FLAG-HiBiT*_fw_PciI for pKF060 |
| prKF258 | ATGAGAGATTTATATTCAGTGACTGCTTTTCTATGTGATGAAACAGAGACAG | expression *FLAG-HiBiT*_rv_SalI for pKF059 |
| prKF261 | AAAAAGAGTCACTGAATATGCATCTCTCATGATATAAGTCGACGG | Site-directed mutagenesis of *byeA*$_{K305A}$_fw |
| prKF267 | GAGCCGGTCGACTTATATCATGAGAGATGCATATTCAG | Site-directed mutagenesis of *byeA*$_{K305A}$_rv_SalI |
| prKF275 | ATCCAACTTCCTTTGTCATCGTCATCCTTG | expression *HiBiT-FLAG*_rv for pKF062 |
| prKF276 | ACGATGACAAAGGAAGTTGGATCAAAGCC | expression of *byeA*$_{64\text{-}309}$_fw for pKF062 |
| prKF279 | GAGCCGGTCGACTTACGGTTTAAGTAAAGACTTATATTCAG | expression of *avrA*$_{Sal}$_rv_SalI for pKF063 |
| prKF287 | AGGTATTATTGCAAAATTAGCACAGGACATTGCTGATGG | Site-directed mutagenesis of *byeA*$_{E58A}$_fw |
| prKF288 | TGTCCTGTGCTAATTTTGCAATAATACCTTTTAATTG | Site-directed mutagenesis of *byeA*$_{E58A}$_rv |
| prMO001 | GCGGGATCCAAGAAGGAGATATACAAATGGTGAGC | expression *HiBiT-FLAG*_fw_BamHI for pMO006 |
| prMO010 | GATTTTTTTTTTTGTCATCGTCATCCTTGTAATC | expression *HiBiT-FLAG*_rv for pMO006 |
| prMO011 | GACGATGACAAAAAAAAAAATCGACCATCCCCTC | expression *bepD*$_{Bhe}$_fw for pMO006 |
| prMO012 | GCGGGTACCTTACATACCAAAGGCCATTC | expression *bepD*$_{Bhe}$_rv_KpnI for pMO006 |

**Table S4.** Accession numbers of the sequences used in this study

| Species | Strain | NCBI Accession Number |
|---|---|---|
| *Bartonella alsatica* | CIP 105477 | NZ_CP058235 |
| *Bartonella clarridgeiae* | 73 | NC_014932 |
| *Bartonella elizabethae* | NCTC12898 | NZ_LR134527 |
| *Bartonella grahamii* | as4aup | CP001562 |
| *Bartonella quintana* | Toulouse | NC_005955 |
| *Bartonella sp.* | *Hoopa Fox 11B* | Ga0127389_11 |
| | *1-1C* | NZ_CP019489 |
| | *CDC skunk* | Ga0117958_11 |
| | *Coyote* 22sub2 | Ga0114331_11 |
| | *JB-15* | NZ_CP019787 |
| | *Racoon 60* | Ga0114332_11 |
| *Bartonella taylorii* | IBS296 | CP083444 |
| *Bartonella tribocorum* | CIP 105476 | NC_010161 |
